# Widespread phages exhibit depth-structured infection coupled with ammonia oxidation

**DOI:** 10.64898/2026.06.18.733297

**Authors:** Yiting Qin, Hao Li, Dinesh Kumar Kuppa Baskaran, Alice Turnham, Maureen Coleman, Karthik Anantharaman, LinXing Chen

## Abstract

Ammonia oxidation is a rate-limiting step in the nitrogen cycle, yet viral contributions to this process remain largely unresolved. Here, we identify three genomically distinct groups of *amoC*-encoding phages (155–338 kilobases in length; termed as amoC-phages) from multiple freshwater lakes in Europe and North America, including the Laurentian Great Lakes. These phages are highly divergent in phylogeny, genome architecture, and gene content, and are predicted to infect two distinct Nitrosomonadaceae ammonia-oxidizing bacterial lineages. The placement of phage-encoded *amoC* genes across these divergent viral clades indicates independent acquisition of *amoC*. Time-series and depth-resolved metagenomes and metatranscriptomes reveal persistent and depth-structured distributions of amoC-phages and their predicted hosts, with seasonal mixing periodically reshaping their co-occurrence patterns. Furthermore, virome data from Lake Mendota show that some of the amoC-phages occur as free viral particles, supporting active viral lysis and particle redistribution along the water column. Metatranscriptomes of the Laurentian Great Lakes reveal coordinated expression of phage structural genes (e.g., major capsid protein) together with phage-encoded *amoC*, indicating active infection *in situ*. Together, these results support a framework in which amoC-phage infection is depth-structured, seasonally dynamic, and coupled to ammonia-oxidizing bacterial host activity, highlighting viruses as previously overlooked components of freshwater nitrogen cycling.

## Introduction

Ammonia oxidation is a key step in the biogeochemical cycling of nitrogen ^1^, linking reduced nitrogen pools to the downstream nitrification process and influencing nutrient availability and primary productivity in various ecosystems, as well as eutrophication and algal blooms in aquatic ecosystems ^2,3^. This process is catalyzed by ammonia monooxygenase, a membrane-bound enzyme complex encoded by the *amoCAB* operon in ammonia-oxidizing bacteria (AOB), including members of *Nitrosospira*, *Nitrosomonas*, and uncultivated lineages such as VFJL01 and BJGV01 ^4–6^, and ammonia-oxidizing archaea (AOA) of Thaumarchaeota ^7^. Among the three subunits, *amoA* is widely used as a functional marker gene to explore the diversity and distribution of AOB and AOA ^8^, while *amoC* represents a conserved core component of the enzyme complex, making it a particularly significant target for viral metabolic manipulation.

Viruses are the most abundant biological entities on Earth ^9–11^. Bacteriophages (phages) are viruses that infect bacteria exclusively and can reshape microbial metabolism during infection by encoding auxiliary metabolic genes (AMGs), thereby altering host fitness and biogeochemical function ^12,13^. Such metabolic reprogramming has been documented in many key pathways, for example, photosynthesis ^14,15^, sulfur metabolism ^16^, and methane oxidation ^17–19^, to name a few, highlighting viruses as active participants in ecosystem processes rather than passive agents of mortality ^13^. Archaeal viruses carrying the *amoC* gene have been reported from marine environments in multiple studies ^20–22^, and more recently, the first amoC-encoding phage was identified in the Eastern Tropical South Pacific oxygen minimum zone, with its amoC most closely related to that of *Nitrosomonas* ^23^. Together, these findings raise the compelling possibility that phages may directly modulate ammonia oxidation in nature, potentially influencing one of the most biogeochemically consequential microbial processes on Earth. However, the ecological distribution, host range, and evolutionary diversity of *amoC*-encoding phages remain poorly resolved. These questions are particularly relevant in freshwater ecosystems. Compared with the oceans, freshwater lakes and reservoirs often exhibit pronounced vertical heterogeneity and seasonal dynamics, including thermal stratification, hypolimnetic oxygen depletion, and episodic water-column mixing ^24–26^. These features strongly structure oxygen and nutrient gradients, thereby shaping the niches, activity, and vertical distributions of ammonia-oxidizing microorganisms ^5,6^. Because ammonia oxidation is a key, and often rate-limiting, step in freshwater nitrogen cycling, *amoC*-encoding phages that infect ammonia oxidizers and potentially modulate their metabolism during infection could represent an overlooked component of freshwater nutrient dynamics. However, whether *amoC*-encoding phages are widespread in freshwater ecosystems, which ammonia oxidizers they infect, how stable their populations are through time, and whether their distributions track environmental gradients and host community structure remain unknown.

Here, we identify large phages encoding *amoC* from multiple geographically distinct freshwater lakes located in Europe and North America. Genome-resolved analyses and host prediction indicate that these phages are primarily associated with the uncultivated Nitrosomonadaceae AOB lineages VFJL01 and BJGV01. Notably, distinct phage groups show contrasting patterns of distribution and persistence. Time-series metagenomic analyses further reveal marked seasonal and depth-related dynamics, including evidence for long-term persistence of some populations and elevated microdiversity in others. Together, these results expand the known diversity and habitat range of *amoC*-encoding phages and suggest that viruses may represent an overlooked component of nitrification ecology in freshwater ecosystems.

## Results

### Discovery of three distinct groups of large phages encoding amoC subunits in freshwater lakes

Reanalysis of the Lake Mendota metagenomic samples spanning 20 years ^18^ identified one virus-like sequence containing an *amoC* gene (Fig. 1a). The *amoC* gene was most similar to those from VFJL01, a genus related to *Nitrosospira* and *Nitrosomonas*. Manual curation and genome extension of this *amoC*-encoding sequence obtained a complete genome (Fig. 1a, Extended Data Fig. 1a), which had a length of 163,870 bp and encoded hallmark viral structural and packaging genes consistent with tailed dsDNA viruses within Caudoviricetes, supporting its classification as a phage (Extended Data Fig. 1b). Hereafter, we refer to such *amoC*-encoding viruses as amoC-phages.

**Fig. 1.**
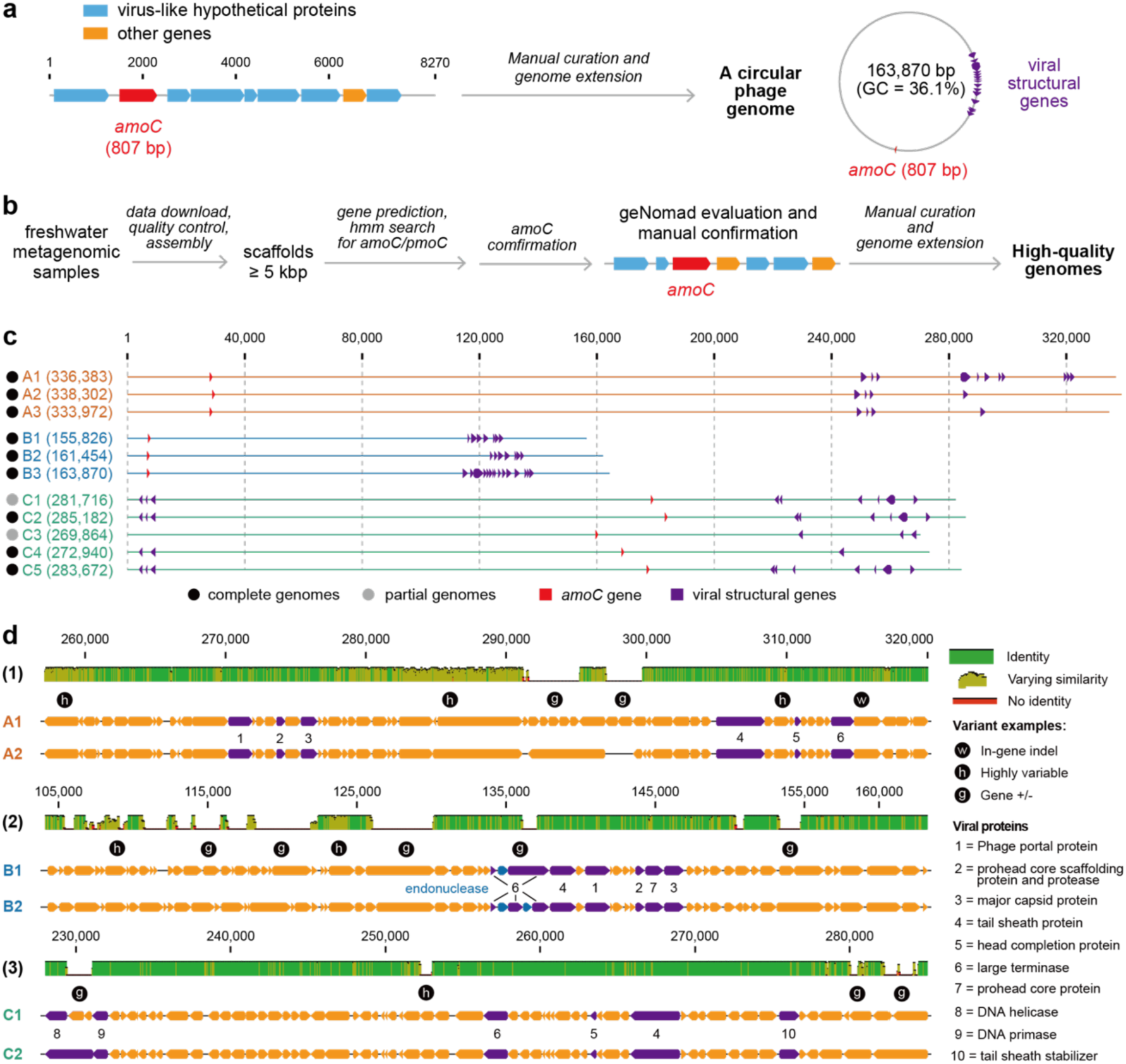
Identification of amoC-phages in freshwater metagenomic samples. (a) A Lake Mendota scaffold containing an *amoC* gene and several virus-like genes was manually curated into a complete phage genome. (b) A pipeline to identify amoC-phage genomes in global freshwater lake metagenomic datasets. The *amoC* and *pmoC* genes are homologous and share the same HMM, and the candidates were confirmed to be *amoC* genes from well-known ammonia-oxidizing bacteria (AOB) and archaea (AOA) using a BLASTp search. The virus-like genes on the *amoC*-encoding scaffolds were determined using an MMseqs2 search against UniProtKB references. (c) The genomic architecture of the 11 curated high-quality amoC-phage genomes. The genomes were assigned to three different groups based on genome-wide average nucleotide identity. The locations of the *amoC* gene (in red) and viral structural genes (in purple) are shown on the genomes. (d) Genomic comparison of within-group members to show their divergence. Comparison of three pairs is shown here, including (1) A1 and A2, and (2) B1 and B2, which are all from Rimov Reservoir, and (3) C1 and C2, with C1 from Lake Michigan and C2 from Lake Superior. Structural and DNA metabolism-related genes are shown in purple, with their corresponding annotations shown on the right.

To reveal the potential prevalence of amoC-phages in freshwater ecosystems, we collected and analyzed 1,244 public metagenomic datasets from freshwater lakes and reservoirs (Fig. 1b, Supplementary Table 1). We obtained 11 high-quality genomes, with nine of them being complete, via manual curation and genome extension (Fig. 1c, Table 1). These amoC-phage genomes could be assigned to three groups, within which members shared >95% genome-wide identity. We termed them group A, group B, and group C, respectively. Both group A and group B comprise three complete genomes, with genome sizes of 333–338 kbp and 155-163 kbp, respectively. Group C includes five genomes (three are complete) and exhibits intermediate genome sizes of 269–285 kbp. These genomes had a GC content of 36.1-39.3%, and encoded 227-433 predicted protein-coding genes and 1-6 tRNA genes. Notably, the CheckV-evaluated completeness values are not 100% for the nine complete genomes, likely due to the lack of closely related references in the CheckV database.

**Table 1.** Summary of amoC-phage genomes. All complete genomes are circular and without assembly errors or gaps. The completeness values from CheckV are listed. The protein-coding genes were predicted using Prodigal (-m -p meta). The tRNAs were predicted using tRNAscan-SE. Note that for the complete genomes, the CheckV completeness values, which were AAI-based with a “medium-confidence” level, are generally lower than those we evaluated based on paired-end read mapping.

| Group | Genome ID | Source | Status | CheckV completeness (%) | Taxonomy (vConTACT3) | Genome length (bp) | GC contents (%) | No. of protein-coding genes | No. of tRNAs |
| --- | --- | --- | --- | --- | --- | --- | --- | --- | --- |
| A | A1 | Rimov Reservoir (Czech) | Complete | 95.22 | Arenbergviridae | 336,383 | 37.9 | 432 | 6 |
|  | A2 | Rimov Reservoir (Czech) | Complete | 95.78 | Arenbergviridae | 338,302 | 37.8 | 423 | 5 |
|  | A3 | Lake Malstasjön (Sweden) | Complete | 94.53 | Arenbergviridae | 333,972 | 37.7 | 433 | 6 |
| B | B1 | Rimov Reservoir (Czech) | Complete | 93.43 | Kyanoviridae | 155,826 | 36.1 | 227 | 3 |
|  | B2 | Rimov Reservoir (Czech) | Complete | 97.71 | Kyanoviridae | 161,454 | 36.4 | 235 | 3 |
|  | B3 | Lake Mendota (USA) | Complete | 98.62 | Kyanoviridae | 163,870 | 36.1 | 229 | 4 |
| C | C1 | Lake Michigan (USA) | Partial | 80.77 | Arenbergviridae | 281,716 | 39.2 | 340 | 1 |
|  | C2 | Lake Superior (USA) | Complete | 81.77 | Arenbergviridae | 285,182 | 39.2 | 345 | 1 |
|  | C3 | Flathead Lake (USA) | Partial | 77.37 | Arenbergviridae | 269,864 | 39.2 | 338 | 1 |
|  | C4 | Lake Zurich (Switzerland) | Complete | 78.25 | Arenbergviridae | 272,940 | 39.2 | 330 | 1 |
|  | C5 | Lake Lugano (Switzerland/Italy) | Complete | 81.33 | Arenbergviridae | 283,672 | 39.3 | 340 | 1 |

Within-group comparable genomic analyses indicated significant divergence between the amoC-phage genomes (Fig. 1d). The dominant differences involved gene gain or loss and intragenic single-nucleotide polymorphisms that generated highly variable regions. Notably, the large terminase genes of the group B amoC-phages are fragmented into two or three pieces by inserted endonuclease genes.

### Phylogeny and gene content of amoC-phages

vConTACT3 analyses suggest that all three groups belonged to Caudoviricetes, with group B assigned to the family Kyanoviridae, whereas groups A and C to Arenbergviridae. At lower taxonomic ranks, all three groups represented novel subfamilies within their corresponding families (Supplementary Table 2). These assignments were supported by ViPTree analysis, in which groups A and C clustered together, whereas group B was affiliated with well-known cyanobacterial phages within Kyanoviridae, including cyanobacterial phages (Extended Data Fig. 2). Interestingly, the closest ViPTree references for groups A and C were viruses associated with animal gut or sewage environments, including *Escherichia*, *Klebsiella*, and *Pseudomonas* phages.

To compare gene contents beyond sequence-level similarity, we annotated protein-coding genes using ColabFold-based structural prediction and annotation (Methods). Principal coordinates analysis (PCoA) based on the presence and absence of protein structure-cluster profiles separated the freshwater amoC-phages into distinct group-level clusters (Fig. 2a). Groups A and C were positioned closer to each other than to group B. By contrast, group B was placed nearer to cyanobacterial phages, reflecting its broader Kyanoviridae-like gene-content background. The previously reported marine *Nitrosomonas*-associated amoC-phage St17_oxy_54E was positioned in the vicinity of group B and cyanobacterial phage references, but remained separated from the freshwater group B amoC-phages.

**Fig. 2.**
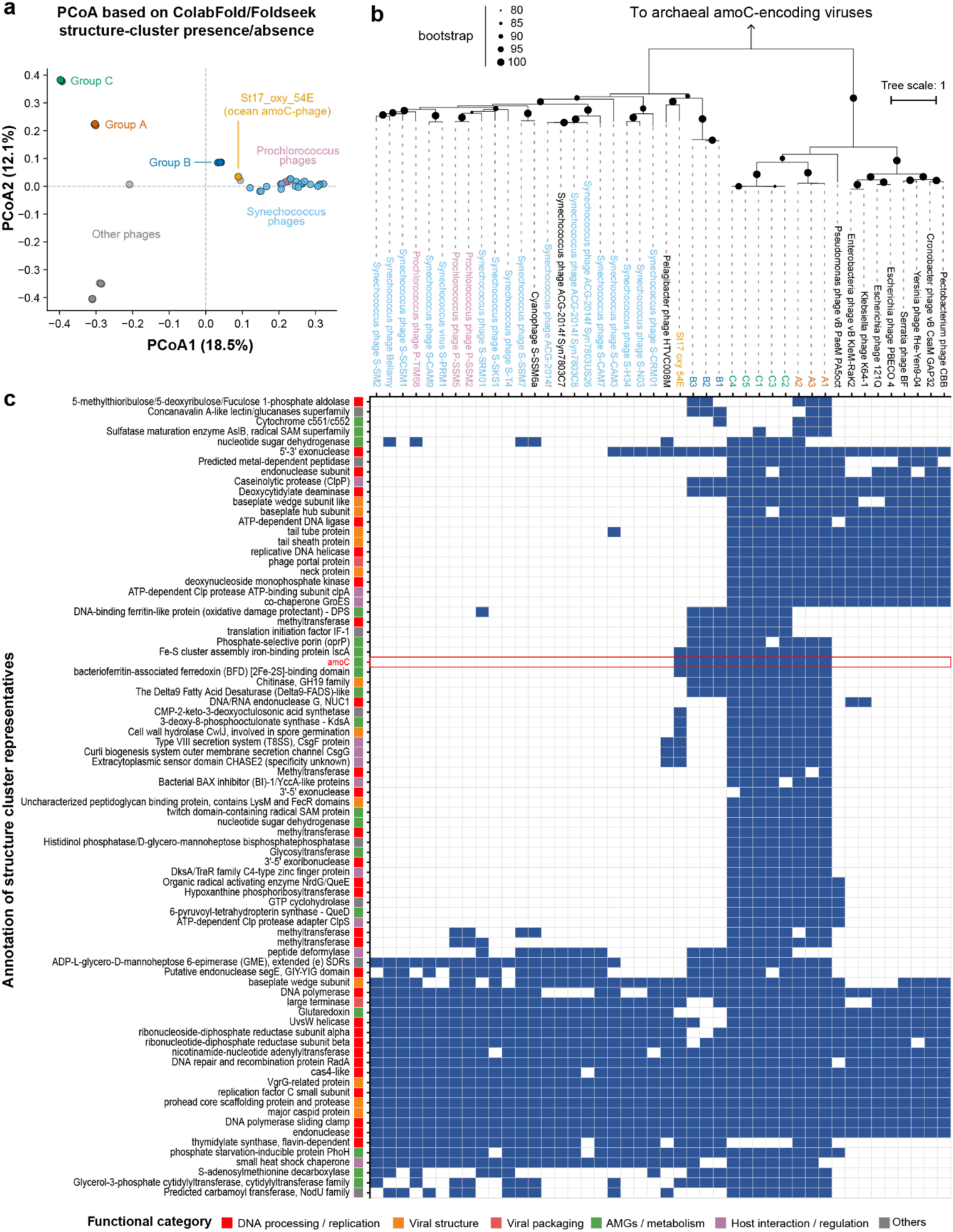
Phylogeny and gene contents comparison of amoC-phages and their relatives. (a) PCoA of amoC-phages and related viruses based on presence-absence profiles of protein structure clusters. Protein structures predicted using ColabFold were clustered using Foldseek to generate a genome-by-structure- cluster count matrix, which was then converted to presence-absence data before calculating pairwise Jaccard distances between genomes. The analysis included the previously reported marine amoC-phage St17_oxy_54E, freshwater amoC-phages identified in this study, and reference viruses selected based on ViPTree analyses. Each point represents one viral genome. Colors indicate viral groups or reference categories. (b) Phylogeny of amoC-phages and related viruses based on concatenated viral marker proteins. The tree was inferred from the concatenated alignment of large terminase, major capsid protein, and phage portal protein sequences, in this order. The tree was rerooted using *amoC*-encoding thaumarchaeal viruses as the outgroup. (c) Presence-absence profiles and functional annotations of protein structure clusters shared by at least two freshwater amoC-phage groups. Rows represent protein structure clusters and were hierarchically clustered using average linkage based on Jaccard distances calculated from binary presence-absence profiles. Columns represent viral genomes and are shown in the same order as in the phylogenetic tree. Blue and white cells indicate presence and absence, respectively. The color strip denotes broad functional categories assigned to each protein structure cluster, with their annotations shown on the left.

We next constructed a phylogenetic tree using concatenated viral marker proteins from all amoC-phages and ViPTree-retrieved reference viruses, together with marine *amoC*-encoding Thaumarchaeota viruses (Methods). The phylogeny showed that the three freshwater amoC-phage groups formed distinct lineages (Fig. 2b). Notably, St17_oxy_54E was placed near group B in the tree. However, the low amino acid identity of the large terminase protein between St17_oxy_54E and group B amoC-phages (∼42%) and the limited genome-wide synteny between them indicated substantial genomic divergence (Extended Data Fig. 3).

To further resolve the gene content of the freshwater amoC-phages, we focused on the annotated protein structure clusters detected in at least two of the three phage groups (Fig. 2c, Supplementary Table 3). Clustering of these shared structure clusters based on their presence–absence matrix revealed a modular gene-content pattern. Groups A and C retained a broad set of shared structure clusters, whereas group B showed a more divergent and reduced profile across many of these clusters. St17_oxy_54E shared only a subset of the structure clusters detected in freshwater amoC-phages and did not recapitulate the full structure-cluster profile of any freshwater group, further supporting its distant relationship to group B inferred from marker-gene phylogeny and whole-genome comparison.

### AmoC-phages are predicted to infect freshwater AOB in the VFJL01 or BJGV01 lineages

To predict the hosts of amoC-phages, we first compared phage-encoded AMGs with their bacterial homologs. The *amoC* genes encoded by groups A and B were most closely related to VFJL01 homologs, with paired protein identities of 93–94%. By contrast, group C phage AmoC proteins shared 85–94% and 98–99% amino acid identity with homologs from marine and freshwater BJGV01, respectively. Phylogeny of phage- and bacteria-encoded AmoC further supported these host assignments and suggested independent acquisition of *amoC* across multiple viral lineages (Fig. 3a, Extended Data Fig. 4). Consistently, predicted structures of phage-encoded AmoC closely matched those of their inferred host homologs, with nearly identical six-transmembrane-helix architectures and low structural deviations between paired phage–host proteins (Extended Data Fig. 5). Analyses of OprP and three additional group C AMGs, SufB, HpnH, and a NapC/NirT-family multiheme cytochrome c, further showed that these phage-encoded proteins grouped with homologs from their predicted freshwater hosts (Extended Data Fig. 6, Supplementary Figs. 1-3). Collectively, these AMG-based analyses support VFJL01 and BJGV01 as the likely bacterial hosts of amoC-phages, consistent with independent host predictions from iPHoP (Supplementary Table 4) and local CRISPR-Cas spacer targeting analyses (Extended Data Fig. 7).

**Fig. 3.**
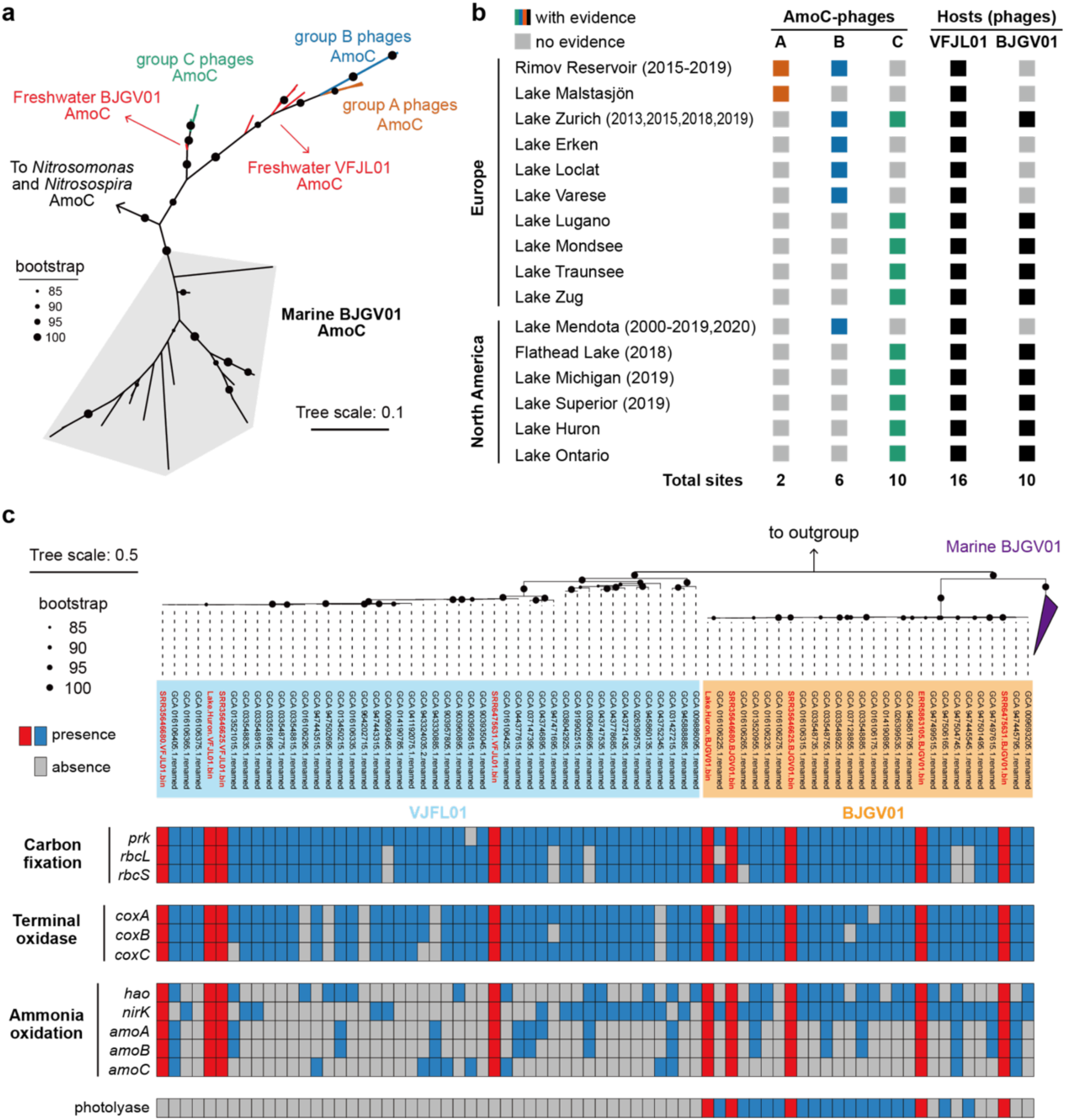
Freshwater AOB lineages VFJL01 and BJGV01 as bacterial hosts of amoC-encoding phages. (a) The phylogeny of phage-encoded and bacterial *amoC* genes. The tree was based on the protein sequences of the *amoC* genes. The *amoC* sequences from both public VFJL01 and BJGV01 MAGs and those reconstructed in this study are included. The details of *Nitrosomonas* and *Nitrosospira* are not shown. For details, refer to the full phylogenetic tree in Extended Data Fig. 4. (b) The identification of amoC-phages and their predicted hosts in freshwater lakes. The detection of amoC-phages was based on metagenomic genome or genomic fragments, and confirmed by paired-end reads mapping. The presence of the hosts was based on the reconstructed VFJL01 and BJGV01 MAGs or their ribosomal protein S3 (rpS3) in the assembled metagenomic datasets. The sampling years of the lakes with data shown in Figs. 4-6 are shown in the brackets. (c) The phylogeny and metabolisms of VFJL01 and BJGV01. The phylogeny was based on the concatenated protein sequences of 120 bacterial marker genes. The genomes reconstructed in this study are highlighted in red. Note that we failed to reconstruct a VFJL01 MAG from Lake Malstasjön as the assembly was fragmented due to low sequencing coverage. prk, phosphoribulokinase; rbcL, ribulose-bisphosphate carboxylase large chain; rbcS, ribulose-bisphosphate carboxylase small chain; coxA/coxB/coxC, cytochrome c oxidase subunit I/II/III; HAO, hydroxylamine dehydrogenase; nirK, nitrite reductase (NO-forming); amoA/amoB/amoC, ammonia monooxygenase subunit A/B/C.

The predicted hosts of amoC-phages, VFJL01 and BJGV01, are two recently defined AOB genus-level lineages and have been reported in freshwater ecosystems ^5,6^. To date, a total of 42 VFJL01 metagenome-assembled genomes (MAGs) and 64 BJGV01 MAGs have been reported (Supplementary Table 5). However, no isolate has been reported. Notably, many of the publicly available freshwater VFJL01 and BJGV01 MAGs were obtained from the same lakes where amoC-phages were detected (Fig. 3b). We noticed that only a small subset of the freshwater VFJL01 and BJGV01 MAGs contain *amoC* genes (Fig. 3c), which is likely due to assembly fragmentation using short paired-end reads ^27^. We additionally reconstructed 4 VFJL01 and 5 BJGV01 MAGs from the metagenomic datasets we assembled in this study (Supplementary Table 6), and manual genome curation was performed to retrieve their *amoC* genes (see details in Supplementary Information). These 9 newly reconstructed MAGs have an average completeness of 86% and an average contamination of 0.9% via CheckM2 evaluation. A phylogenetic tree of VFJL01 and BJGV01 indicated their relatedness, with BJGV01 from marine habitats clustered separately from freshwater ones (Fig. 3c). Functional analyses of all the freshwater MAGs revealed their metabolic potentials in carbon fixation and ammonia oxidation.

We identified 10 freshwater lakes from Europe and 6 freshwater lakes in North America, with at least one amoC-phage group and their corresponding predicted bacterial hosts (Fig. 3b). Interestingly, group C amoC-phages were recovered from all the Laurentian Great Lakes, except Lake Erie. Notably, although most lakes contained both VFJL01 and BJGV01, amoC-phages identified in this study that linked to both host genera were only observed in Lake Zurich.

### Groups A and B amoC-phages exhibit distinct population dynamics despite long-term persistence in the Rimov Reservoir

To characterize the spatial and temporal dynamics of VFJL01-associated amoC-phages, we analyzed long-term metagenomic samples collected from 0.5 m and 30 m in Rimov Reservoir ^28,29^, where both groups A and B amoC-phages were detected. Across multiple years (2015–2019), amoC-phages and their predicted VFJL01 hosts were consistently enriched in deep waters at 30 m, with group A amoC-phages generally having higher abundance (Fig. 4a, Supplementary Table 7). This depth restriction was transiently disrupted during seasonal mixing (typically starting in November), when both phages and hosts became detectable at both depths, often with elevated abundance of phages in surface waters (0.5 m). In contrast, during the spring bloom period, group B amoC-phages were largely undetectable despite the continued presence of their hosts.

**Fig. 4.**
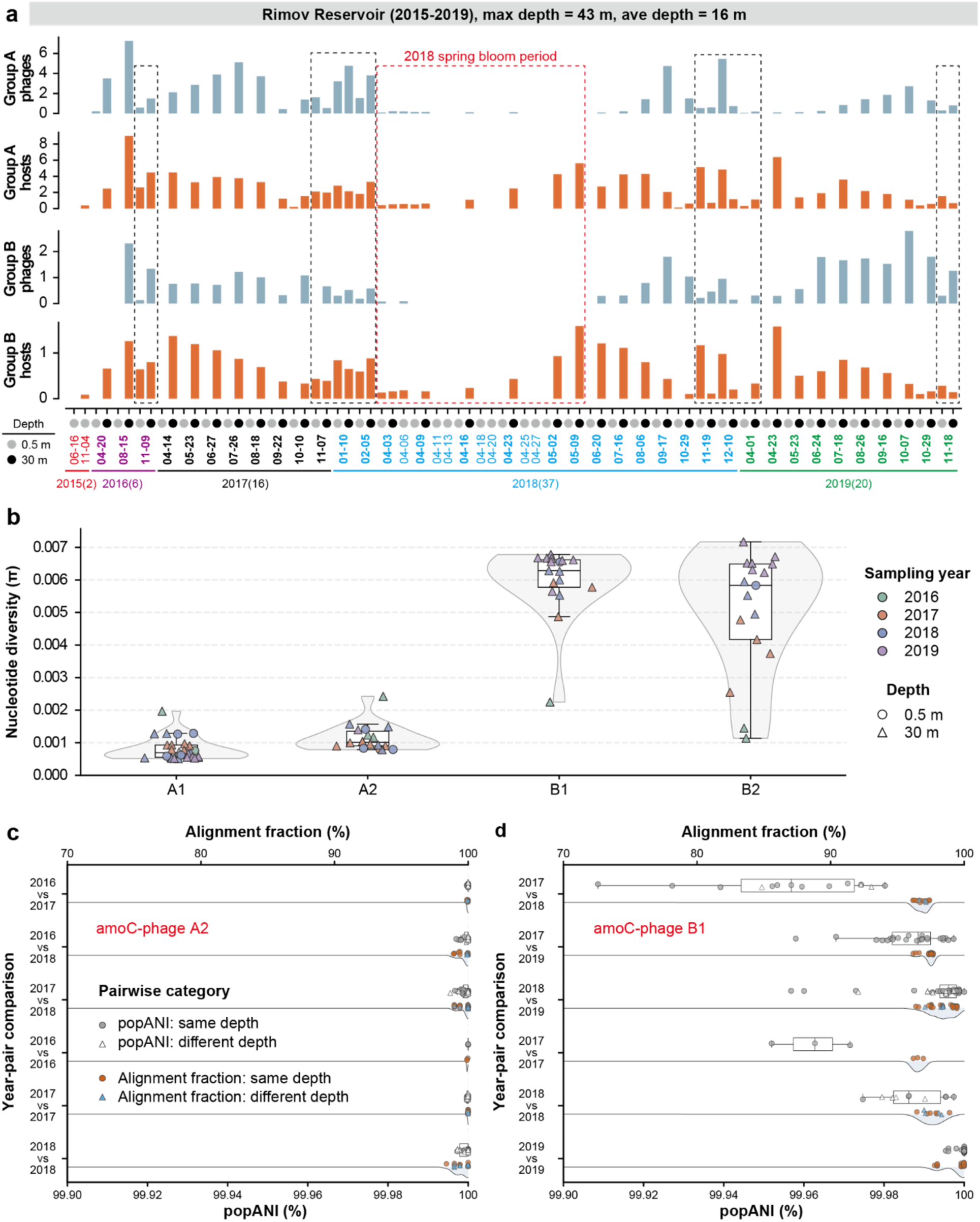
Population dynamics and structure of groups A and B amoC-phages in Rimov Reservoir. (a) Temporal and depth-resolved abundance (RPKM) of group A (A1, A2) and group B (B1, B2) amoC-phages and their predicted VFJL01 hosts from 2015 to 2019. Samples were collected at 0.5 m and 30 m. Phages and hosts are consistently enriched in deep waters and transiently detected in surface waters during seasonal mixing. The dashed red box indicates the 2018 spring bloom period, when group B amoC-phages were largely undetectable despite host presence. The dashed black boxes indicate the sampling points with amoC-phages detected at both depths. (b) Within-sample nucleotide diversity (π) of amoC-phage populations across years. Violin plots show the distribution of nucleotide diversity values for each phage group, with boxplots and individual samples overlaid. Group A phages consistently exhibit low nucleotide diversity and narrow distributions, whereas group B phages display substantially higher and more variable diversity across samples and years. Samples collected from 0.5 m and 30 m are indicated by circles and triangles, respectively. (c-d) Pairwise population comparisons for amoC-phage A2 (c) and amoC-phage B1 (d). Lower x-axes indicate pairwise population average nucleotide identity (popANI), whereas upper x-axes indicate pairwise alignment fraction. Boxplots summarize popANI distributions across year-pair comparisons, and ridge plots show the corresponding alignment fraction distributions. Individual comparisons are overlaid and categorized by whether the paired samples originated from the same depth or different depths.

To determine whether amoC-phage populations redistributed across the water column remained genetically stable through time, we quantified within-sample nucleotide diversity and pairwise between-sample population similarity using inStrain ^30^ (**Methods**). Group A amoC-phages consistently exhibited low nucleotide diversity (π) across samples, indicating genetically homogeneous populations with limited microdiversity (Fig. 4b). Pairwise sample comparisons further showed that amoC-phages A1 and A2 maintained highly similar populations across years, with consistently high popANI and near-complete alignment fractions in both within-year and between-year comparisons (Fig. 4c, Extended Data Fig. 8a).

In contrast, group B amoC-phages displayed substantially higher nucleotide diversity (π) and broader diversity distributions across samples and years (Fig. 4b). Pairwise sample comparisons of B1 and B2 revealed pronounced temporal structuring despite consistently high alignment fractions (Fig. 4d, Extended Data Fig. 8b). In particular, populations associated with B1 in 2019 formed a highly homogeneous cluster that was differentiated from populations detected in earlier years (Fig. 4d), whereas populations associated with B2 remained highly similar between 2016 and 2017 but became progressively differentiated from populations detected in 2018 and 2019 (Extended Data Fig. 8b). These patterns are consistent with temporal shifts in dominant closely related populations rather than gradual diversification within a single stable population.

### Group B amoC-phages persisted over two decades and were likely active near the oxycline in Lake Mendota

Building on the persistent but temporally structured group B amoC-phage populations observed in Rimov Reservoir, we next examined whether related phages also persist in a second freshwater system and whether their distributions are structured by seasonal mixing and depth. We focused on Lake Mendota, which has a maximum depth of 25.3 m and a mean depth of 12.8 m, and where group B amoC-phages and their predicted VFJL01 hosts were detected (Fig. 3b).

Analysis of the samples collected in 2000–2019 from the upper mixed layer of Lake Mendota (integrated to 12 m depth, generally above the oxycline) ^18^ revealed recurrent detection of both group B amoC-phages and VFJL01 hosts across years (Fig. 5a). Notably, phage and host abundances increased markedly during the Ice-on period (Fig. 5a, Supplementary Table 8). These Ice-on samples were characterized by lower temperature, relatively higher dissolved oxygen, and reduced Secchi depth compared with open-water seasonal phases (Extended Data Fig. 9). Because these samples represent integrated upper-water communities, this enrichment likely reflects seasonal redistribution and retention of VFJL01 populations and associated amoC-phages in the upper water column following mixing and under-ice stabilization. Reduced light exposure under ice may favor VFJL01 persistence, particularly because the reconstructed VFJL01 genomes lack recognizable UV-damage repair genes (Fig. 3c), although other under-ice conditions, including oxygen availability and ammonia regenerated or redistributed during mixing, may also contribute.

**Fig. 5.**
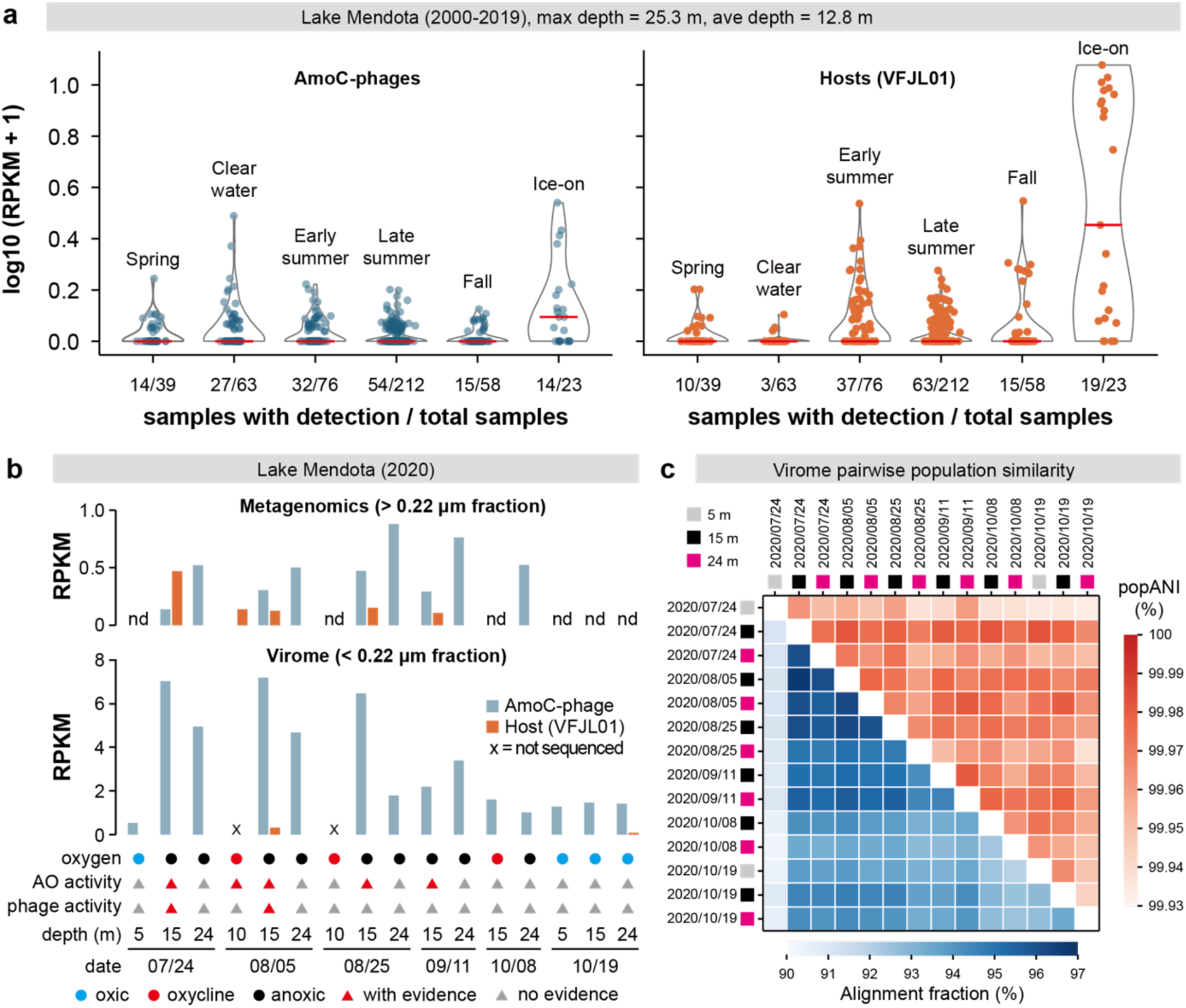
Distribution and persistence of group B amoC-phages in Lake Mendota. (a) Co-occurrence of group B amoC-phages and their predicted hosts in Lake Mendota over 20 years (2000–2019). Each point represents a composite sample from the upper 12 m, grouped by seasonal phase. Red lines indicate median values. Note that RPKM values are scaled for visualization. nd, not detected. (b) Comparative metagenomic, virome, and metatranscriptomic profiles of group B amoC-phages and their predicted hosts in Lake Mendota, based on samples collected in summer 2020. (c) Pairwise population similarity of group B amoC-phages based on virome samples collected during summer 2020. Upper triangles show popANI values, and lower triangles show alignment fractions calculated using inStrain. Samples are ordered by sampling date and depth. Pairwise comparisons revealed consistently high popANI values together with high alignment fractions across depths and sampling dates, indicating that group B amoC-phages detected throughout the water column belonged to closely related and genetically well-connected populations rather than depth-specific divergent lineages.

To resolve the depth structure underlying these long-term upper-water patterns, we further analyzed meta-omic samples collected in summer 2020 from 10 m, 15 m, and 24 m in Lake Mendota ^31^. Group B amoC-phage DNA was frequently enriched in the 24 m water samples, whereas VFJL01 hosts were primarily detected at 10-15 m (Fig. 5b, Supplementary Table 9). These host-positive depths corresponded to the oxycline-to-upper-anoxic transition during summer stratification, and metatranscriptomic signals for host ammonia oxidation were likewise concentrated at 10-15 m as well. By contrast, the amoC-phage signal in the virome fraction often peaked at 15 m rather than 24 m. Detectable but limited phage transcriptional signals were observed in two 15 m samples (Extended Data Fig. 10). These depth-resolved patterns suggest that host activity and infection- or lysis-associated viral signals were concentrated near the oxycline and upper anoxic boundary, whereas deeper waters may retain or accumulate phage-derived DNA and/or particles.

We next asked whether the recurrently detected amoC-phages represented related or genetically divergent phages. Because phage abundance in the 20-year metagenomic time series was generally low, pairwise between-sample popANI comparisons were restricted to samples with sufficient sequencing depth (13 samples in total). The recurrently detected group B amoC-phage populations remained closely related across years (popANI values ≥99.94%; Supplementary Table 10), supporting long-term persistence of related viral populations rather than repeated detection of unrelated phages. We further performed the same analyses using the virome samples collected in summer 2020 (Fig. 5c). The near-identical popANI values (≥99.93%), together with high alignment fractions (93.7% on average), indicate that the detected viral signals originated from closely related and genetically well-connected populations rather than from depth- or date-specific divergent lineages. The absence of strong genetic partitioning across depths is consistent with the redistribution or exchange of group B amoC-phage populations throughout the water column. However, the high but incomplete alignment fractions indicate that these populations were not strictly clonal and retained measurable genomic heterogeneity across depths and sampling dates.

### Group C amoC-phages are widespread and transcriptionally active in the Laurentian Great Lakes

Group C amoC-phages represent the most broadly distributed lineage identified in this study (Fig. 3b). Across multiple freshwater systems spanning Europe and North America, including Lake Zurich and Flathead Lake, group C amoC-phages and their predicted BJGV01 hosts were consistently co-detected, with a general tendency toward higher abundance in deeper waters, where host signals were often strongest, though they were also intermittently detected in shallower layers (Figs. 6a and b).

**Fig. 6.**
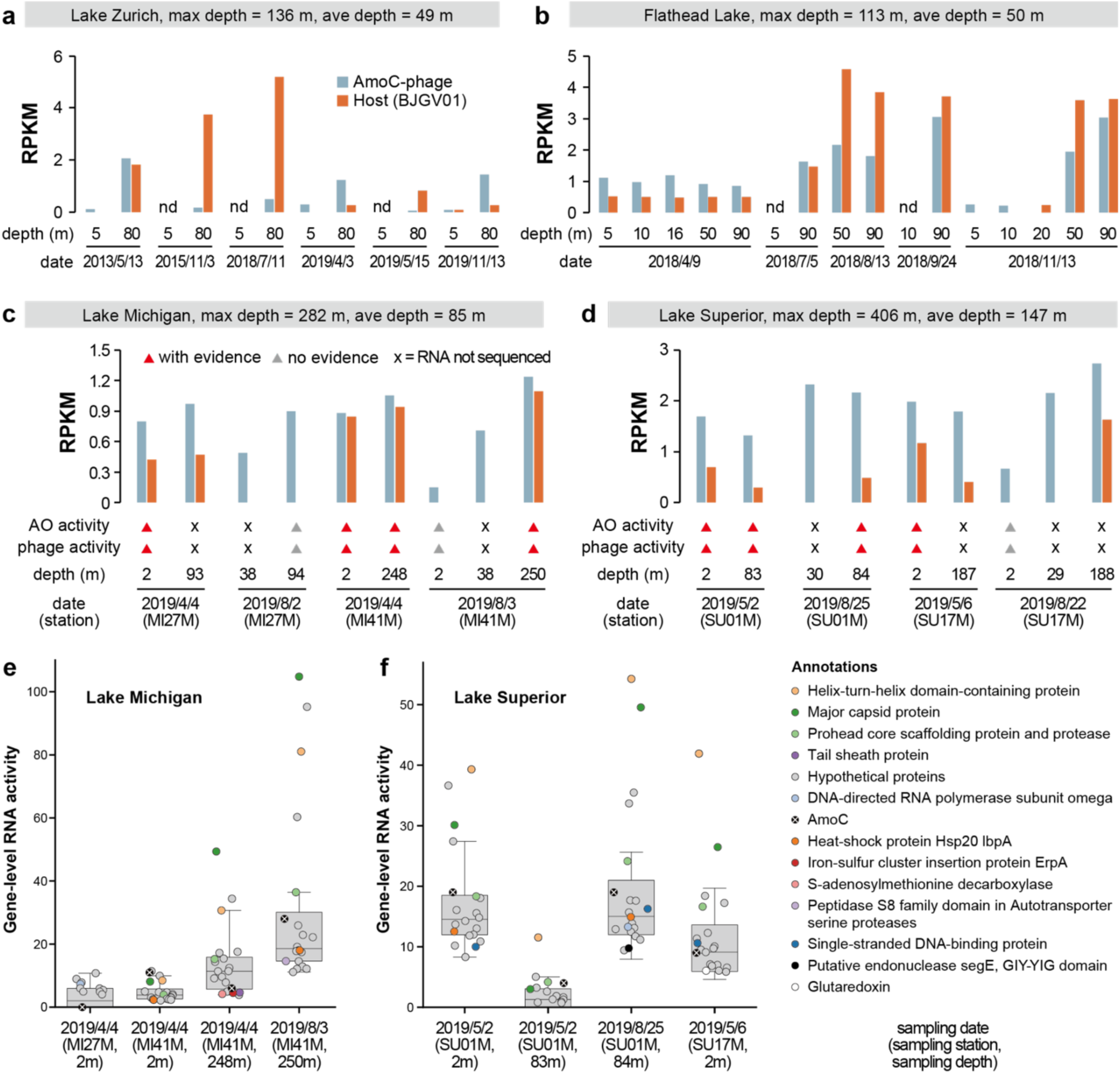
The distribution and transcriptional activity of group C amoC-phages. The detection of group C amoC-phages and their hosts in (a) Lake Zurich (Switzerland) samples collected from 2013 to 2019 and (b) Flathead Lake (USA) samples collected in 2018. The presence and transcriptional activities of amoC-phages and the hosts in (c) Lake Michigan and (d) Lake Superior samples collected in 2019. Lake Michigan and Lake Superior belong to the Laurentian Great Lakes in North America. In (c) and (d), the information on the sampling dates and stations is shown at the bottom. (e, f) The transcriptional activities of genes encoded by group C amoC-phages in (e) Lake Michigan and (f) Lake Superior. For each sample, only the 20 genes with the highest transcriptional activity are shown; if fewer than 20 genes were transcriptionally detected, all detected genes are shown. The Boxplots summarize the distribution of gene transcriptional activity across samples, with individual genes overlaid as points.

Group C amoC-phages were also detected in the Laurentian Great Lakes, including samples collected in 2012 and 2019 from Lake Michigan and Lake Superior, and samples of Lake Ontario and Lake Huron as well (Supplementary Tables 11 and 12). However, we did not identify any known amoC-phages in Lake Erie, which also belongs to the Laurentian Great Lakes.

In Lake Michigan and Lake Superior, both group C amoC-phages and BJGV01 hosts were detected across a broad depth range at multiple sampling stations (Figs. 6c and d, Supplementary Fig. 4). Notably, during spring sampling in April and May, both phages and hosts were frequently detected in shallow waters, in several cases together with evidence for ammonia-oxidizer activity and phage transcriptional activity. Metatranscriptomic analyses further revealed that group C amoC-phages expressed a specific subset of genes *in situ* (Figs. 6e and f). These genes were predominantly involved in virion assembly and morphogenesis, including the major capsid protein, the prohead core scaffolding protein and protease. In contrast, genes involved in DNA packaging, such as the portal protein and large terminase, showed no detectable expression. Importantly, transcripts of phage-encoded *amoC* were also detected, consistent with active expression of this AMG during infection. The observed transcriptional profile is indicative of late-stage infection activity, suggesting ongoing or recent virion production rather than early stages of infection. In this context, phage-encoded *amoC* expression may occur during the late phase of infection, potentially contributing to host ammonia oxidation capacity at a stage when cellular metabolism is being redirected toward virion production.

## Discussion

Ammonia oxidation is the first and often rate-limiting step of nitrification, a central process in the nitrogen cycle ^1,32^, yet current conceptual models of freshwater nitrification largely treat microbial activity as decoupled from viral processes ^33–35^. Here, we identify *amoC*-encoding phages (i.e., amoC-phages) as widespread, persistent, and, in some cases, transcriptionally active members of nitrifier-associated communities (Figs. 4-6). These phages are phylogenetically diverse and associated with two ammonia-oxidizing bacterial lineages (Fig. 3), indicating that viral acquisition of *amoC* has occurred independently across distinct viral groups.

Beyond their diversity, our analyses reveal that amoC-phage dynamics are shaped by lake-specific vertical structure and recurring seasonal mixing (Figs. 4-6). The clearest depth-resolved evidence comes from Lake Mendota, where metabolically active VFJL01 hosts were concentrated near the oxycline, free viral particles peaked at intermediate depths, and phage-derived metagenomic signals were strongest in deeper waters (Fig. 5). This pattern suggests vertical partitioning between the likely zone of host activity and viral production near the oxycline, and deeper waters where viral particles or phage-derived DNA may accumulate or persist. In other lakes, amoC-phages and their predicted hosts were generally more abundant in deeper water layers, whereas seasonal water-column reorganization, including mixing or transitional stratification, periodically redistributed them into shallower waters (Figs. 4 and 6). Thus, seasonal mixing plays a key role in modulating these interactions by redistributing hosts and phages and periodically resetting infection niches. However, because comparable fine-scale physicochemical profiles and matched depth-resolved activity measurements are not available for some of these freshwater systems, we cannot determine whether the active phage-host interaction zones in these lakes are also centered near the oxycline as observed in Lake Mendota. Together, these findings indicate that amoC-phages are unlikely to be incidental carriers of metabolic genes alone but are recurrently associated with ammonia-oxidizing bacterial populations and, in some cases, actively express their *amoC* genes during infection-related activity. These observations support a framework in which amoC-phage activity is spatially and seasonally coupled to ammonia-oxidizing bacterial niches, while the precise depth of active infection may vary among lakes and remains best resolved in systems with matched physicochemical and meta-omic data.

Host physiological and metabolic traits provide a plausible context for interpreting the contrasting distributions of VFJL01- and BJGV01-associated amoC-phages. The BJGV01-associated group C phages were detected across broader depth ranges during spring mixing in Lake Michigan and Lake Superior, including transcriptionally active signals in shallow 2 m samples collected in April and May (Fig. 6). The presence of photolyase genes in the BJGV01 genomes reconstructed from these lakes (Fig. 3c) suggests that these hosts may be better able to tolerate light-associated DNA damage, potentially allowing BJGV01-associated phage–host interactions to persist when host populations are redistributed into illuminated surface waters during mixing. In Lake Mendota, where group B phages are associated with VFJL01, long-term upper-water metagenomes showed elevated phage and host abundances during the ice-on period (Fig. 5). Because light exposure is reduced under ice cover, these observations suggest that low-light winter conditions, together with seasonal mixing or retention of deeper-water populations in the upper water column, may create a favorable window for VFJL01-associated amoC-phage and host co-occurrence. Nutrient regimes may further influence amoC-phage distributions indirectly by selecting for different ammonia-oxidizer lineages. For example, Lake Erie, which is much shallower than Lake Michigan and Lake Superior, harbors distinct nitrifier communities with little evidence for VFJL01 and BJGV01 lineages ^5^, likely reflecting its substantially higher NH4^+^ and nutrient concentrations compared with the other Laurentian Great Lakes ^36^.

The distribution patterns and population genetic structures of the three amoC-phage groups indicate that they represent distinct ecological strategies rather than a single homogeneous entity (Figs. 1 and 2). Group A appears to be a geographically restricted and genetically stable lineage (Fig. 4), suggesting long-term persistence within a narrow ecological niche. Group B exhibits broader distribution, high microdiversity, and strong virome signals, consistent with dynamic lytic populations in which infection and particle production are clearly and closely linked to environmental gradients, such as the oxycline, in Lake Mendota (Fig. 5). Group C is widely distributed across limnologically distinct lakes and shows clear transcriptional activity *in situ* (Fig. 6), indicating an ecologically active lineage capable of sustained infection across diverse conditions.

A significant biological implication of our results is that all three amoC-phage groups encode *amoC* rather than *amoA* or *amoB* (Fig. 2). This pattern is reminiscent of several well-characterized viral AMG systems, in which viruses encode selected components of host metabolic pathways or enzyme complexes rather than complete pathways ^14,17,19–21^. In ammonia-oxidizing microorganisms, additional standalone copies of *amoC* are frequently observed and have been proposed to support enzyme stability or function under environmental stress ^37–39^. The independent acquisition of *amoC* across divergent viral lineages suggests that it represents a particularly advantageous target for viral modulation of host metabolism. Consistent with this interpretation, the *amoC* of group C phage is transcriptionally active along with structural genes (Fig. 6), indicating that it is expressed during active infection. This raises the possibility that viral *amoC* supplements or stabilizes host ammonia monooxygenase function during late-stage infection, potentially maintaining ammonia oxidation capacity while cellular resources are redirected toward virion production.

Together, our findings demonstrate that amoC-carrying phages are widespread, persistent, and functionally active components of freshwater ecosystems. By revealing that freshwater phages encoding key nitrification genes are embedded within depth-structured and seasonally dynamic ecological interactions, this study expands current views of nitrogen cycling to include virus-mediated metabolic modulation as an integral component in freshwater lake ecosystems. More broadly, these results suggest that viral contributions to biogeochemical processes may be more pervasive and mechanistically structured than previously appreciated, particularly in environments characterized by strong spatial and temporal heterogeneity.

## Methods

### Sample collection in the Laurentian Great Lakes

Water samples from the Laurentian Great Lakes were collected aboard the U.S. EPA vessel *R/V Lake Guardian*. Full details are described in Hernandez Limon and Coleman (under revision). Briefly, samples were collected from several stations in Lake Michigan and Lake Superior during 2019, including periods of spring mixing (April/May) and summer stratification (August). For each station, water samples were collected from selected depths for DNA/RNA sequencing using a conductivity-temperature-depth (CTD) rosette sampler (Sea-Bird Scientific). Corresponding water chemistry analyses, along with water column profiles of temperature, dissolved oxygen, and fluorescence, were measured by the EPA as part of the Great Lakes Water Quality Survey program. These environmental metadata are freely available via the EPA Central Data Exchange portal (https://cdx.epa.gov). Sample metadata, including location, depth, and date, are provided in Supplementary Table 11.

For metagenomic and metatranscriptomic analyses, lake water was prefiltered through a GF/A glass fiber filter (Whatman 1820-047; nominal pore size 1.6 μm), and picoplankton cells were collected on 0.22 μm polyethersulfone filters (Millipore Express GPWP). Filters for DNA and RNA extraction were immediately flash frozen in liquid nitrogen and stored at -80 °C until processing. DNA was extracted using a modified phenol-chloroform extraction protocol ^40^. RNA was extracted using the mirVana miRNA Isolation kit (Invitrogen), following the manufacturer’s instructions. RNA extracts were treated with DNase (Turbo DNA-free, Invitrogen) to remove residual DNA, and RNA quality was assessed using Bioanalyzer. Ribosomal RNA was depleted using QIAseq FastSelect (Qiagen), and metatranscriptomic libraries were prepared from the remaining RNA. Sequencing was performed on an Illumina NovaSeq S4 platform by the Joint Genome Institute (Walnut Creek, CA, USA) to generate paired-end reads of 150 bp in length.

### Retrieval and analysis of freshwater metagenomic samples

Public freshwater metagenomic datasets were retrieved from the NCBI Sequence Read Archive and publications, using keywords and metadata associated with lakes, reservoirs, ponds, and freshwater microbial communities, and only those sequenced with paired-end short reads were included. Metagenomic datasets from freshwater lakes and reservoirs, including Lake Mendota, Rimov Reservoir, and the Laurentian Great Lakes, were retrieved. Sample metadata were listed in Supplementary Table 1. The raw sequencing metagenomic reads were downloaded from NCBI and quality-filtered using fastp version 1.0.1 with default parameters ^41^ to remove sequencing adapters and low-quality bases and reads. The quality reads were assembled for each sample individually using metaSPAdes version 3.15.5 with the k-mer set of “21,33,55,77,99,127” ^42^. For some lakes, as the abundance of amoC-phages was low, we thus co-assembled all the related paired-end reads using metaSPAdes. The assembled scaffolds shorter than 5 kbp were excluded from viral discovery analyses, unless they were used for manual curation and genome extension of amoC-containing scaffolds. The protein-coding genes were predicted from assembled scaffolds using Prodigal version 2.6.3 (-m -p meta) ^43^.

### Identification of *amoC*-encoding viral scaffolds

To identify candidate *amoC*-encoding viral sequences, predicted protein-coding genes from assembled scaffolds were searched against amoC/pmoC hidden Markov models (K10946 from Kofam ^44^) using hmmsearch from HMMER version 3.4 ^45^, with a score threshold of 133.93 (which is suggested in Kofam). Because amoC and pmoC genes can be detected by the same HMM profile, all candidate amoC/pmoC proteins were further compared against well-known bacterial AmoC and PmoC reference sequences retrieved from GTDB r226 genomes ^46^ using BLASTp ^47^. Candidate genes were retained as amoC only when their best matches and phylogenetic placement were consistent with amoC from AOB rather than pmoC from bacterial methanotrophs.

Those identified scaffolds encoding *amoC* genes were then evaluated for viral origin. The amoC-containing scaffolds were evaluated using geNomad version 1.11.1 ^48^, and VirSorter2 version 2.2.4 ^49^ with default parameters, and their protein-coding genes were also searched against UniProtKB ^50^ using MMseqs2 version 15.6f452 ^51^. Viral hallmark genes, including genes encoding major capsid protein, phage portal protein, large and small terminase, tail, baseplate, and phage DNA replication proteins, were evaluated by a combination of the UniProtKB-based sequence similarity searches, geNomad and VirSorter2 prediction, and manual confirmation using online BLASTp (https://blast.ncbi.nlm.nih.gov/BlastAlign.cgi). Scaffolds were considered candidate amoC-phage scaffolds when they encoded amoC together with one or more viral hallmark genes and/or lacked sufficient evidence for being bacterial chromosomal or plasmid fragments.

### Manual curation and genome extension

The purpose of manual curation of the candidate *amoC*-encoding viral scaffolds was to exclude potential assembly errors, extend incomplete scaffolds, close gaps, and recover complete amoC-phage genomes where possible. For each *amoC*-encoding viral scaffold to be curated, we mapped the corresponding quality paired-end reads using Bowtie2 version 2.5.4 with default parameters ^52^. The mapping profiles were filtered to exclude those paired-end reads that were not mapped using shrinksam (available at https://github.com/bcthomas/shrinksam), and the SAM file was sorted and converted to a BAM file with SAMtools version 1.22.1 using default parameters ^53^. The *amoC*-encoding scaffold under curation and the corresponding BAM file were imported into Geneious Prime release 2025.1.2 ^54^, and the paired-end reads were remapped to the scaffold using the “Map to Reference(s)” function (with the mapper of “Geneious”) in Geneious Prime, with “Custom Sensitivity” allowing no mismatch.

We first identified the regions on the scaffolds with no reads mapped, which suggested assembly errors or nucleotide variations; these regions were annotated as “gap” in Geneious Prime. We also flagged the regions with obviously lower coverage than those of neighboring regions, that usually because of nucleotide variations, and such regions were annotated as “check” in Geneious Prime. We then manually checked the two end regions to see if reads were mapped across the ends. Once detected, we obtained the consensus sequences of the overhang reads and added them to the ends of the scaffold, so the scaffold was extended, usually tens of base pairs long.

Second, we mapped the same paired-end reads as used above to the *amoC*-encoding viral scaffold that we annotated, flagged, and extended in Geneious Prime using the “Map to Reference(s)” function (with the mapper of “Geneious”) with “Medium Sensitivity/Fast”. We did not use “Custom Sensitivity” and “no mismatch” this time, as we needed to confirm the “gap” and “check” regions with all available paired-end reads. If the annotated or flagged regions contained nucleotide variations, we curated the scaffold using the variable reads with higher abundance. However, if the annotated regions have assembly errors, we fixed them as we described previously ^27^, and the procedures are available here (https://banfield-lab.gitbook.io/ggkbase-help-center/Z7AVfDAiORm7FGbuC8xs/genome-curation/scaffold-extension-and-gap-closing). We then checked the two ends for overhang reads and used their consensus sequences to extend the scaffold. Hereafter, we reran the “Map to Reference(s)” function with “Medium Sensitivity/Fast” using the unmapped reads from the last run of mapping and performed the fixation and extension multiple times until no more reads could be mapped. Note that the assembly errors may not be able to be fixed, usually due to insufficient sequencing depth or the need for one or more runs of mapping to obtain more available reads.

Third, we searched the extended fragments against all the assembled scaffolds from the corresponding sample using local BLASTn; the hits that had >95% nucleotide identity and comparative sequencing depth with the *amoC*-encoding viral scaffold were retrieved and assembled with the curated *amoC*-encoding viral scaffold using the “De Novo Assemble” function in Geneious Prime using the assembler of “Geneious” with “Medium Sensitivity / Fast”, and obtained a curated and extended scaffold.

After that, we used Bowtie2 to map all quality paired-end reads to the curated and extended scaffold and performed all the steps as described above in Geneious Prime to fix any unsolved assembly errors and extend the *amoC*-encoding scaffold. Note that these steps may have to be performed many times until the scaffold is curated to a circular genome or no more reads can be retrieved for extension. A basic mechanism to determine a circular genome is that there is sufficient sequence overlap (usually > 50 bp) between the two ends of the scaffold. To confirm the circularization of a genome, we need to exclude the existence of multiple copy genes, such as transposase, at the two ends of the scaffold. The curated genomes were classified as complete when they were circular, lacked detectable gaps or unsupported joins, and showed consistent read-mapping support across the whole genome. Genomes that could not be circularized but contained long viral scaffolds with amoC and multiple viral hallmark genes were retained as partial amoC-phage genomes.

Genome completeness was also evaluated using CheckV version 1.0.3 with the “end_to_end” program ^55^. The CheckV estimates were interpreted cautiously because closely related reference genomes of amoC-phages were largely absent from the CheckV database. Therefore, for complete genomes, manual curation, circular genome structure, and paired-end read support were considered more informative than CheckV completeness alone.

A tool that we developed previously, named COBRA ^56^, is able to perform automatic genome extension similarly. However, it will fail if the viral population is highly variable. We suggest attempting COBRA first to curate any viral genomes of interest, and manual curation and genome extension should be applied if it does not work well.

Note that some specific situations, including the regions encoding *amoC*, phosphate-selective porin gene (*oprP*), the cysteine desulfurase activator complex subunit SufB gene, the hopanoid biosynthesis-associated radical SAM protein HpnH gene, and NapC/NirT-family multiheme cytochrome c gene, that should be carefully evaluated in curating the group C amoC-phage genomes, were included in the Supplementary Information.

### Viral genome clustering, taxonomic classification, and whole-genome comparison

All the curated amoC-phage genomes were clustered into species-level viral groups using the rapid genome clustering approach provided by CheckV (https://bitbucket.org/berkeleylab/checkv/src/master/), with the parameters set as “min_ani = 95, min_tcov = 85”. The taxonomic placement of amoC-phages was assessed using vConTACT3 based on genome-wide protein-cluster profiles ^57^. The resulting classification was used to assign amoC-phages to established viral taxa where possible and to identify novel taxonomic groups. Whole-genome proteomic similarity was further examined using ViPTree ^58^, and the closest reference viruses were used to contextualize the evolutionary placement of each amoC-phage group (see the “**Phylogenetic analysis of amoC-phages**” section below). Genome-wide synteny between selected amoC-phages from the same groups and related references, including the only reported amoC-phage (from the marine environment), St17_oxy_54E, was examined and visualized as genome comparison plots using the “progressiveMauve algorithm” ^59,60^ function in Geneious Prime.

### Host prediction of amoC-phages

We used multiple approaches to predict the bacterial hosts of the amoC-phages. First, we compared the *amoC* genes encoded by phages against all the *amoC* genes from GTDB r226 genomes. Second, we analyzed all 11 curated amoC-phage genomes using iPHoP version 1.3.3 with default parameters ^61^. Third, we predicted the repeat regions using PILER-CR version 1.06 ^62^ from all the scaffolds ≥ 5000 bp assembled from metagenomic data of lakes with amoC-phages detected. The CRISPR spacers were extracted from repeat-containing scaffolds with cas gene(s) identified within 1000 bp of the repeat region, and searched against the curated amoC-phage genomes using blastn-short with an e-value threshold of 1e-4, allowing no more than one mismatch across the whole spacer length. The taxonomic assignment of the scaffolds with targeting spacers was evaluated via the corresponding MAGs (if the scaffolds were binned) and/or the taxonomy of the closest references of the protein-coding genes on the scaffolds. The cas genes were determined by hmmsearch from HMMER version 3.4 ^45^ against the Cas protein-related HMM databases from TIGRFAM ^63^.

### Genome binning and manual curation of host genomes

To reconstruct bacterial hosts associated with amoC-phages, metagenomic assemblies from amoC-phage-containing samples were subjected to genome binning. The quality paired-end reads were mapped to all the assembled scaffolds using Bowtie2 version 2.5.4 with default parameters ^52^, and the scaffold coverage profiles were calculated using the jgi_summarize_bam_contig_depths function from metaBAT2 ^64^. The scaffolds with a minimum length of 2,500 bp were binned using MetaBAT2 with default parameters to obtain metagenome-assembled genomes (MAGs).

Taxonomic classification of the MAGs was performed using GTDB-Tk version 2.4.1 ^65^ with the GTDB r226 genomes as references. The MAGs assigned to Nitrosomonadaceae, particularly VFJL01 and BJGV01, were manually inspected and refined. To recover fragmented ammonia oxidation genes and ribosomal protein S3 scaffolds, contigs associated with VFJL01 and BJGV01 were further examined using coverage patterns, tetranucleotide composition, taxonomic annotation, and linkage to high-confidence MAG scaffolds. Where *amoC*, *amoA*, *amoB*, *hao*, *nirK*, *rbcL*, *rbcS*, *prk*, photolyase, and terminal oxidase genes were fragmented or absent from initial MAGs, local searches of all assembled scaffolds against those sequences from public VFJL01 and BJGV01 genomes were used to identify additional host-derived scaffolds. Only scaffolds with consistent taxonomic affiliation and coverage patterns were assigned to the corresponding VFJL01 or BJGV01 MAGs and included in host metabolic analyses. The refined MAGs were evaluated using CheckM2 version 0.1.3 ^66^ for completeness and contamination.

### Retrieval of public VFJL01 and BJGV01 genomes

The information on the public VFJL01 and BJGV01 genomes was retrieved from GTDB, and the genomes were downloaded manually from NCBI. The relevant information was included in Supplementary Table 5. So far, all the VFJL01 and BJGV01 genomes have been reconstructed from metagenomics. The genome quality of public VFJL01 and BJGV01 genomes was evaluated using CheckM2 version 0.1.3 ^66^, and their taxonomic assignment was confirmed using GTDB-Tk version 2.4.1^65^.

### Gene prediction and annotation of amoC-phage and host genomes

Protein-coding genes were predicted from curated amoC-phage genomes and VFJL01 and BJGV01 MAGs using Prodigal version 2.6.3 (-m -p meta) ^43^. Transfer RNA genes of the amoC-phage genomes were predicted using tRNAscan-SE version 1.4 ^67^. To compare gene contents at the protein-structure level, protein sequences encoded by amoC-phages, marine *amoC*-encoding viruses, and selected reference viruses based on ViPTree analyses were subjected to structure prediction using ColabFold ^68^ downloaded on 22 November 2024, with the parameters set as: “--num-recycle 3, --use-gpu-relax, --amber, --stop-at-score 70”. The predicted structures with a pLDDT score of ≥70 were searched against several reference protein structure databases, including Protein Data Bank (PDB) ^69^, BFVD ^70^, and NCBIFAM ^71^, using FoldSeek 9.427df8a with the “easy-search” command (-c = 0.5, --cov-mode = 0) ^72^. Only those hits with a TM score ≥0.5 (“alntmscore”) were retained for further analyses. The predicted proteins of the host genomes were annotated using hmmsearch against the KOfam hmm database ^44^. The HMM hits were filtered based on the score threshold for each KO provided in the “ko_list” file along with the KOfam HMM database.

### Identification and visualization of shared amoC-phage protein structure clusters

To profile the presence and abundance of phage proteins across all the virus genomes, the predicted protein structures with a pLDDT score of ≥70 were clustered using FoldSeek 9.427df8a with the “easy-cluster” command (--threads 10 -c 0.7 --cov-mode 0 --tmscore-threshold 0.7) ^72^. To identify structure clusters shared among freshwater amoC-phages and the references, a protein structure cluster was retained if it was detected in at least two of the three amoC-phage groups, where detection was defined as the presence of at least one protein structure from any genome within that group. The presence and absence of the retained protein structure clusters were converted to a matrix, and those with functional annotations were clustered using Jaccard distance and average-linkage hierarchical clustering and visualized (Fig. 2c). Functional annotations of each protein structure cluster were based on comparison against reference structure databases as mentioned above.

### Phylogenetic analysis of amoC-phages

Protein-coding genes of amoC-phage genomes and reference viral genomes were predicted or collected from genome annotations. Three conserved viral marker proteins, including the large terminase subunit, major capsid protein, and phage portal protein, were used for phylogenetic reconstruction. For each marker gene, homologous proteins were identified from the amoC-phage genomes and selected reference viruses. Protein sequences were aligned individually using MAFFT, and poorly aligned regions were removed using trimAl. The trimmed alignments of the three marker proteins were concatenated for each genome. A maximum-likelihood phylogenetic tree was then inferred from the concatenated alignment using IQ-TREE2 version 2.4.0 ^73^, with the best-fitting substitution model selected automatically and branch support assessed using ultrafast bootstrap approximation. The resulting tree was visualized and annotated using iTOL ^74^.

### Phylogenetic analysis of VFJL01 and BJGV01

For all the VFJL01 and BJGV01 genomes we reconstructed in this study and downloaded from NCBI, their taxonomy was first evaluated using GTDB-Tk version 2.4.1 ^65^, as mentioned above. A phylogenomic tree was constructed using the concatenated alignment of 120 conserved bacterial marker proteins, which was generated via the GTDB-Tk analysis. A maximum-likelihood tree was inferred using IQ-TREE2 version 2.4.0 ^73^ with automatic model selection and branch support estimated using ultrafast bootstrap approximation. The concatenated alignment of 120 conserved bacterial marker proteins of 10 randomly selected GTDB Alphaproteobacteria genomes was included as outgroups. The phylogenetic tree was visualized and annotated using iTOL ^74^.

### Phage and host abundance calculation

The detection and abundance of amoC-phages and their predicted hosts (VFJL01 and BJGV01) were estimated by read recruitment. For amoC-phages, the reference sequences were the manually curated amoC-phage genomes reconstructed from the corresponding lakes or reservoirs in this study. For the predicted bacterial hosts, the reference sequences were ribosomal protein S3 (rpS3)-encoding scaffolds assigned to VFJL01 or BJGV01. The rpS3 is a single-copy gene and is used as a conservative bacterial marker; thus, the RPKM (see below) calculated for the rpS3-encoding scaffolds could roughly reflect that of the corresponding genomes.

For each sample, the quality paired-end reads were mapped to the corresponding phage genomes or host rpS3-encoding scaffolds using CoverM version 0.7.0 ^75^. The presence of a given amoC-phage genome or host scaffold in a sample was first evaluated using the trimmed mean coverage and minimum covered fraction (≥70%) calculated by CoverM. The read recruitment parameters using CoverM were set as “--min-read-aligned-percent 90 --min-read-percent-identity 90 --min-covered-fraction 70 --contig-end-exclusion 75 -m trimmed_mean”. A phage genome or host rpS3-encoding scaffold was considered present in a sample only when the resulting trimmed mean coverage was greater than 1. Otherwise, the corresponding phage or host was treated as not detected in that sample, even if a small number of reads could be recruited to the reference. This two-step procedure was used to reduce spurious low-level read recruitment and to ensure that reported phage and host abundance values were supported by both high-identity read mapping and sufficient reference coverage.

For samples in which a phage genome or host scaffold was considered present, the number of mapped reads assigned to the corresponding reference was calculated using the same read-recruitment parameters. Abundance was then normalized as reads per kilobase per million reads (RPKM). Specifically, RPKM was calculated as the number of reads mapped to a phage genome or host rpS3-encoding scaffold multiplied by 1,000,000,000 and then divided by the reference length in base pairs and the total number of quality reads of the corresponding sample.

### Microdiversity and population structure analyses

To investigate the microdiversity and population structure of amoC-phages in Rimov Reservoir and Lake Mendota, the corresponding metagenomic or viromic reads (when available) were mapped to amoC-phage genomes reconstructed from the corresponding sites, using Bowtie2 version 2.5.4 with default parameters ^52^.

The microdiversity profiling of the groups A and B amoC-phages in Rimov Reservoir was performed using inStrain version 1.10.0 ^30^. For each sample, the inStrain profile was used to calculate genome coverage, breadth, nucleotide diversity (π), and single-nucleotide variant (SNV) frequencies. Only genome–sample pairs satisfying the following criteria were retained for downstream analyses: genome breadth ≥70%, average coverage ≥10×, and mean read pair identity ≥95% when available. Genome breadth was defined as the fraction of genome positions covered by at least one mapped read. The pairwise population comparisons were performed using the inStrain “compare” operation. To avoid underestimation of pairwise genome overlap caused by uneven viral genome coverage, pairwise comparisons were conducted using a relaxed minimum coverage threshold (-c 1), requiring positions to be covered by at least 1× in both samples to be included in comparisons. Pairwise population average nucleotide identity (popANI) and alignment fraction were extracted from the comparison results. Alignment fraction was defined as the fraction of genome positions that were simultaneously covered and comparable between two samples. For temporal population analyses, pairwise comparisons were grouped according to sampling years (e.g., within-year and between-year comparisons). Pairwise popANI and alignment fraction distributions were visualized using boxplots and ridge-density plots. Within-sample nucleotide diversity distributions were visualized using violin plots with overlaid boxplots and individual sample points. All analyses and visualizations were performed in Python using pandas, NumPy, matplotlib, and SciPy.

For the group B amoC-phages in Lake Mendota, as their sequencing depth in the metagenomic datasets of the 20 years (2000-2019) was generally low (<10×), we thus did not calculate the nucleotide diversity (π). Instead, we profiled the pairwise popANI using the metagenomic reads and also the metagenomic and viromic reads from the 2020 summer samples, as described above. And the pairwise comparisons were grouped according to sampling dates and depths.

### Metatranscriptomic analysis

Metatranscriptomic datasets were available for samples from Lake Mendota, Lake Michigan, and Lake Superior, which made it possible to evaluate the *in situ* transcriptional activities of group B amoC-phages in Lake Mendota and the group C amoC-phages in the two Laurentian Great Lakes, as well as their bacterial hosts in the corresponding samples. The paired-end metatranscriptomic reads were quality-filtered using fastp version 1.0.1 with default parameters ^41^ to remove sequencing adapters and low-quality bases and reads. The quality reads were mapped to curated amoC-phage genomes and host MAGs using Bowtie2 version 2.5.4 with default parameters ^52^, and the SAM file was sorted and converted to a BAM file with SAMtools version 1.22.1 using default parameters ^53^.

Gene-level transcriptional activity was calculated from the resulting sorted and indexed BAM files using pysam ^76^. Gene coordinates were obtained from Prodigal protein FASTA headers, which report the start and end positions of each gene on the corresponding genomes and scaffolds. Alignments were not used if they were unmapped, marked as secondary or supplementary alignments, or flagged as PCR/optical duplicates in the BAM file. For each retained alignment, read-to-reference nucleotide identity was estimated from the BAM NM tag, which records the number of mismatches and indels relative to the reference. Specifically, identity was calculated as one minus the number of mismatches/indels divided by the aligned read length. The aligned read length was calculated from the CIGAR string using the parts of the read that were aligned to the reference, including matches, mismatches, and insertions, but excluding soft-clipped bases. Only alignments with estimated nucleotide identity ≥97% were retained. For each gene, RNA coverage was calculated across the aligned reference positions overlapping the gene. Gene-level transcriptional activity was defined as the mean per-base RNA coverage across the full gene region. Genes were considered transcriptionally detected only when ≥70% of their length was covered by at least one retained RNA read.

The transcriptional activity of amoC-phages was inferred when reads mapped to phage genes, particularly structural or morphogenesis-related genes such as major capsid protein, phage portal protein, large terminase, prohead core scaffolding protein, and protease.

To assess the transcriptional activity of group C amoC-phages in Lake Superior and Lake Michigan, special consideration was required for phage genes that are highly similar to homologs in the predicted BJGV01 hosts (Supplementary Fig. 3). These host-like genes include *amoC*, the phosphate-selective porin gene *oprP*, the cysteine desulfurase activator complex subunit SufB gene, the hopanoid biosynthesis-associated radical SAM protein HpnH gene, and the NapC/NirT-family multiheme cytochrome *c* gene. Because reads derived from phage- and host-encoded homologs cannot be reliably distinguished across highly conserved regions, we focused on sequence-divergent regions between the phage and host genes to obtain a more conservative estimate of phage-derived transcription. Among the five host-like genes, this approach was only feasible for *amoC*, which contains a distinct divergent region near the N-terminus with several nucleotide differences between phage- and bacteria-encoded homologs. To evaluate the transcriptional activity of phage-encoded *amoC*, quality-filtered paired-end metatranscriptomic reads were mapped to the group C amoC-phage genomes, and reads overlapping this phage-specific divergent region (without mismatch) were manually counted. The resulting read counts were used as conservative transcriptional values for phage-encoded *amoC*. Because only the reads overlapping sites that distinguish phage amoC from host amoC were counted, these values likely underestimate the true transcriptional activity of phage-encoded *amoC*.

Host ammonia-oxidizer activity was evaluated based on transcription of ammonia oxidation and nitrification-associated genes, including *amoA*, *amoB*, *amoC*, *hao*, *nirK*, and terminal oxidase genes, together with the transcription of host marker genes, such as ribosomal protein genes, where available. Samples with coordinated expression of host ammonia oxidation genes, phage structural genes, and phage-encoded *amoC* were interpreted as evidence for in situ phage activity linked to ammonia-oxidizing bacterial hosts.

## Data availability

The amoC-phage genomes are available at Figshare via https://figshare.com/s/bb351a75b2e543747f5b.

## Acknowledgments

L-X.C. is supported by the Research Program of the University of Science and Technology of China (KY2400000036, KY2400000040, and GG2400007010). We thank the Supercomputing Center at the University of Science and Technology of China for its support of ColabFold analyses.

## Author contributions

L-X.C. and K.A. designed the study. Y.T.Q., H.L., and L-X.C. analyzed the data. Y.T.Q. and L-X.C. drafted the manuscript. A.T. and M.C. collected and sequenced the samples from the Laurentian Great Lakes. Y.T.Q., H.L., D.K.K.B., A.T., M.C., K.A., and L-X.C. revised the manuscript. All authors approved the manuscript.

---

**Extended Data Fig. 1.**
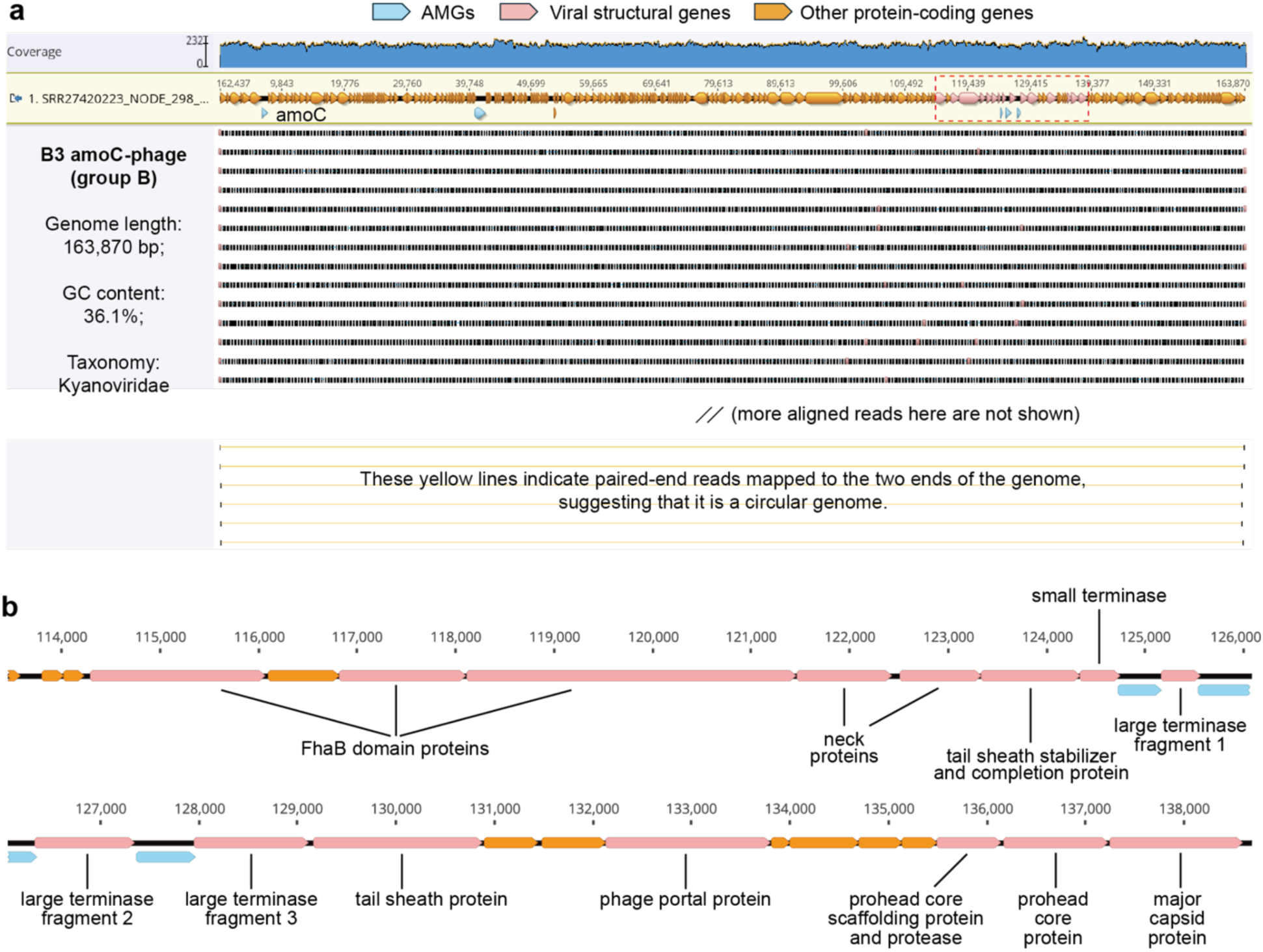
A circular, complete *amoC*-encoding viral genome recovered from Lake Mendota. (a) Read-mapping profile of the B3 amoC-phage genome. Sequencing coverage across the genome is shown at the top, and individual mapped reads are shown below. Paired-end reads spanning the two genome termini are highlighted with yellow links at the bottom, supporting circularization of the assembled genome. The continuous read coverage across the genome, together with the absence of visible coverage breaks or assembly gaps, supports the recovery of a complete circular genome. General genome features, including genome length, GC content, and taxonomic assignment, are shown on the left. Reads were mapped to the genome using Bowtie2 v2.5.4 with default parameters ^52^ and visualized in Geneious Prime ^54^ (release version, 2025.1.2). (b) Expanded view of the region highlighted by the red dashed box in (a), showing hallmark viral structural and DNA-packaging genes, including terminase, portal, prohead, tail sheath, and major capsid genes. Genes are colored by functional categories: auxiliary metabolic genes (AMGs), viral structural genes, and other protein-coding genes.

**Extended Data Fig. 2.**
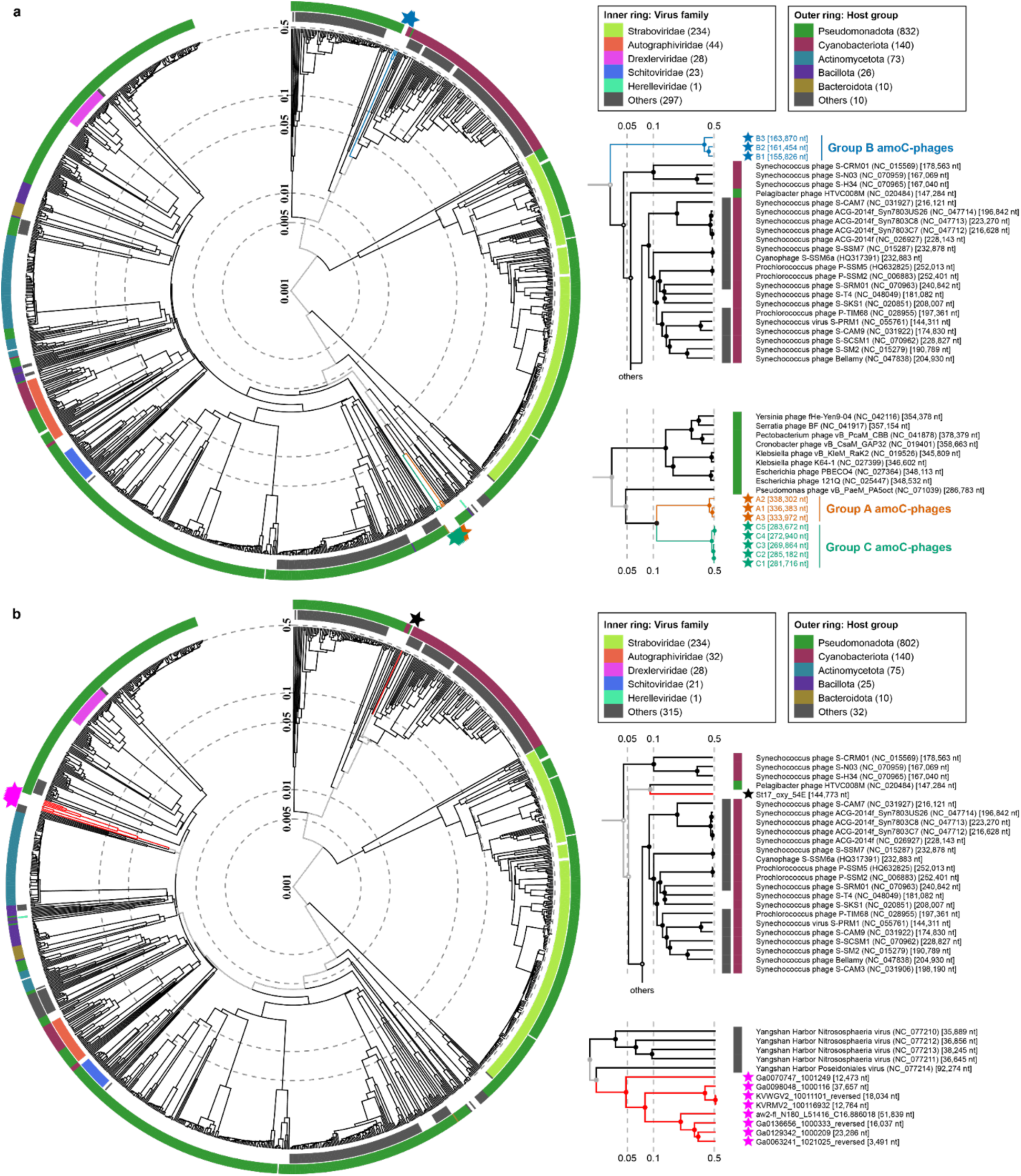
ViPTree-based phylogenomic placement of freshwater and marine *amoC*-encoding viruses. (a) ViPTree-based clustering of freshwater amoC-phages reported in this study, together with representative ViPTree reference viruses. (b) ViPTree-based clustering of previously published marine *amoC*-encoding viruses together with representative ViPTree reference viruses. In both (a) and (b), the inner grey track denotes the family-level taxonomic assignment of reference viruses, and the outer colored track denotes the predicted host group of the corresponding reference viruses. The *amoC*-encoding viruses, including those reported in this study and previously published marine representatives, are indicated by colored stars.

**Extended Data Fig. 3.**
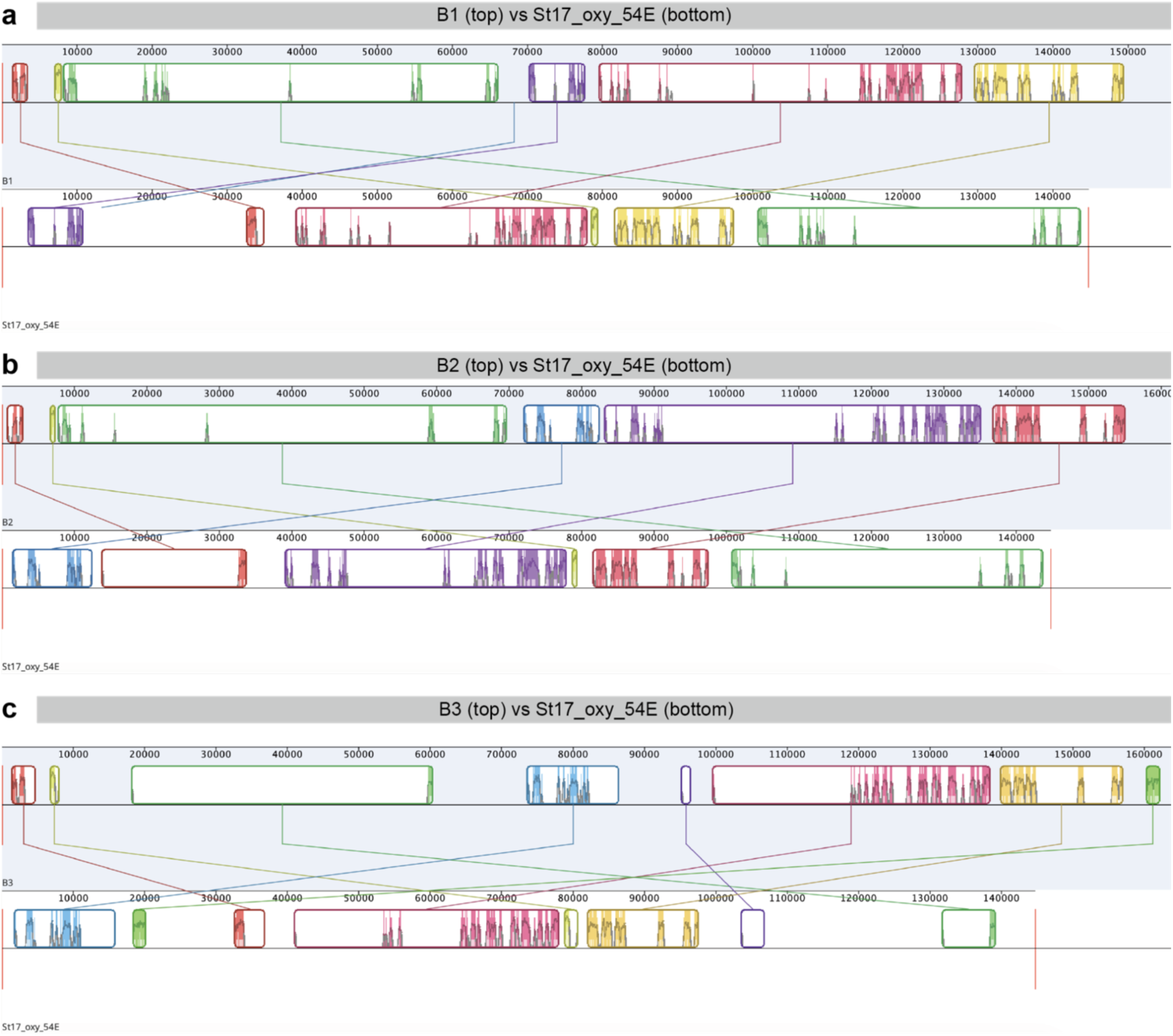
Limited genome-wide synteny between group B freshwater amoC-phages and the marine amoC-phage St17_oxy_54E. Whole-genome comparisons between representative group B freshwater amoC-phages and the previously reported marine *amoC*-encoding virus St17_oxy_54E. The genomes of (a) B1, (b) B2, and (c) B3 are shown on top of each comparison, and St17_oxy_54E is shown at the bottom. Colored blocks indicate locally conserved genomic regions detected between the compared genomes, with connecting lines linking homologous regions. Although St17_oxy_54E was placed near group B amoC-phages in marker-gene phylogenetic analysis, only fragmented and discontinuous regions of similarity were detected across the genomes, and the shared regions showed limited collinearity.

**Extended Data Fig. 4.**
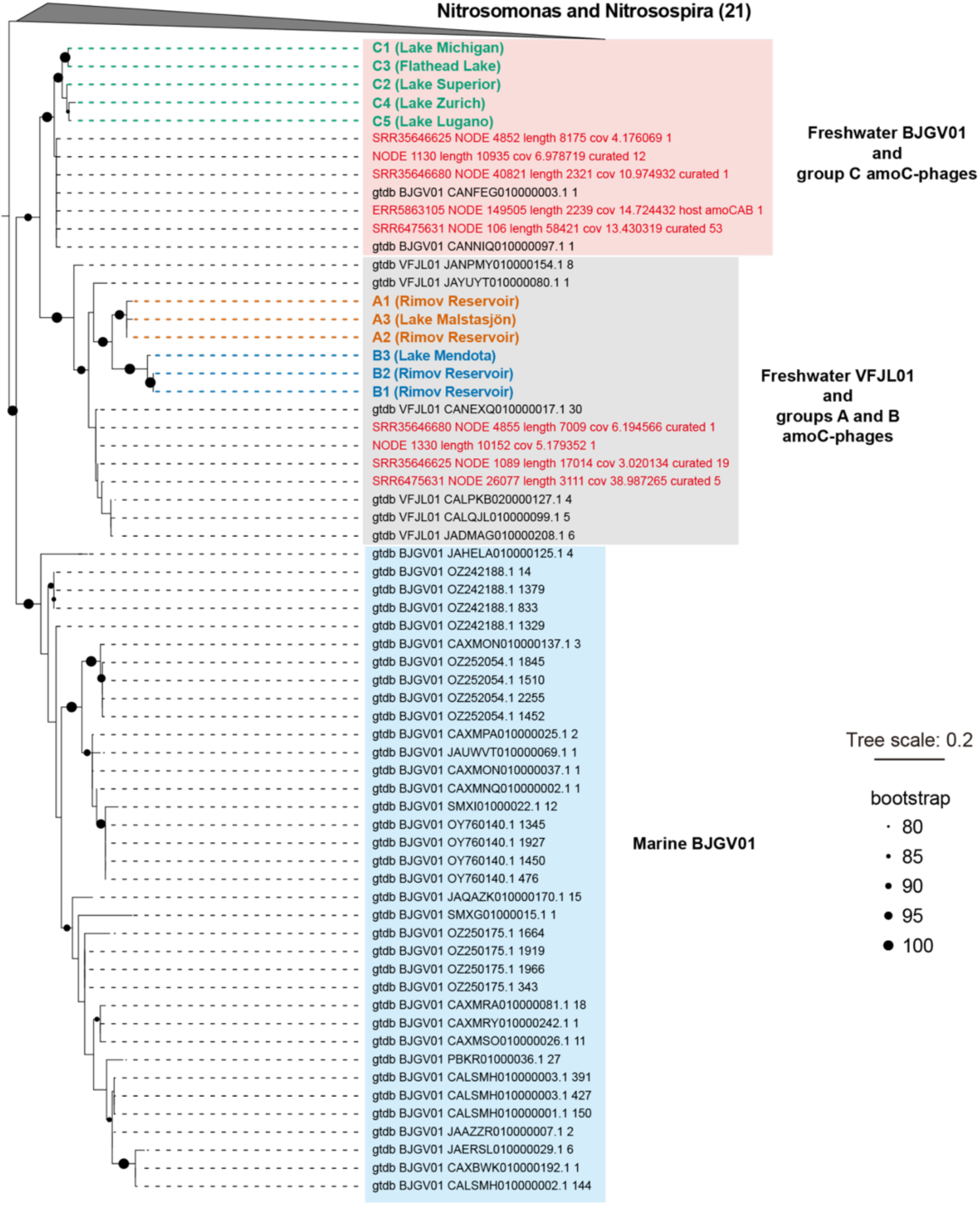
The phylogeny of phage- and bacteria-encoded *amoC* genes. Phylogenetic placement of *amoC* genes encoded by amoC-phages, reconstructed VFJL01 and BJGV01 genomes, and bacterial reference genomes (including *Nitrosomonas* and *Nitrosospira*, and all public VFJL01 and BJGV01 genomes). The AmoC from VFJL01 and BJGV01 genomes reconstructed in this study are labeled in red. The *amoC* genes encoded by amoC-phages are colored according to phage group, and the lakes from which the amoC-phage genomes were reconstructed are shown in parentheses after genome IDs. AmoC sequences from *Nitrosospira* and *Nitrosomonas* reference genomes are collapsed, with the number of included sequences shown in parentheses. Dashed lines indicate the positions of amoC-phage-encoded *oprP* genes. The tree scale bar indicates amino acid substitutions per site.

**Extended Data Fig. 5.**
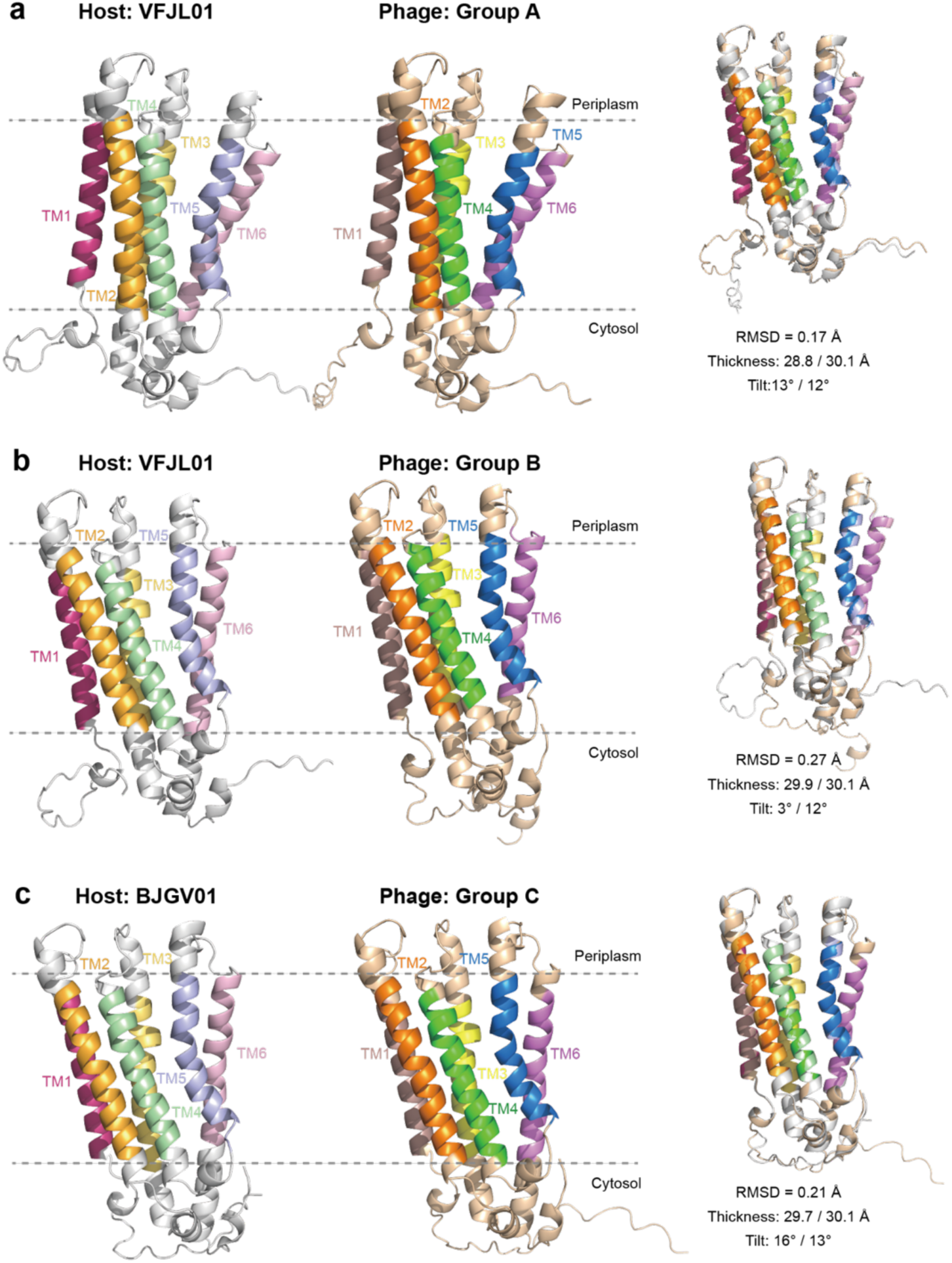
Structural conservation and membrane topology of viral and host AmoC proteins. The AmoC structures of (a) group A amoC-phage and the predicted host, (b) group B amoC-phage and the predicted host, and (c) group C amoC-phage and the predicted host. The representative host and phage AmoC proteins were modeled using AlphaFold2 and structurally superimposed in PyMOL. All viral and host AmoC proteins exhibited highly conserved six-transmembrane-helical architectures (TM1–TM6), with pairwise RMSD values ranging from 0.17 to 0.27 Å, indicating near-identical overall protein folds. Membrane positioning analysis using PPM 3.0 further revealed comparable hydrophobic thicknesses (28.8–30.1 Å) and membrane tilt angles (3–16°) between viral and host proteins, suggesting similar membrane insertion properties and orientations. These results demonstrate that phage-encoded AmoC proteins retain the structural and topological features of their host homologs, supporting their potential incorporation into ammonia monooxygenase complexes during infection.

**Extended Data Fig. 6.**
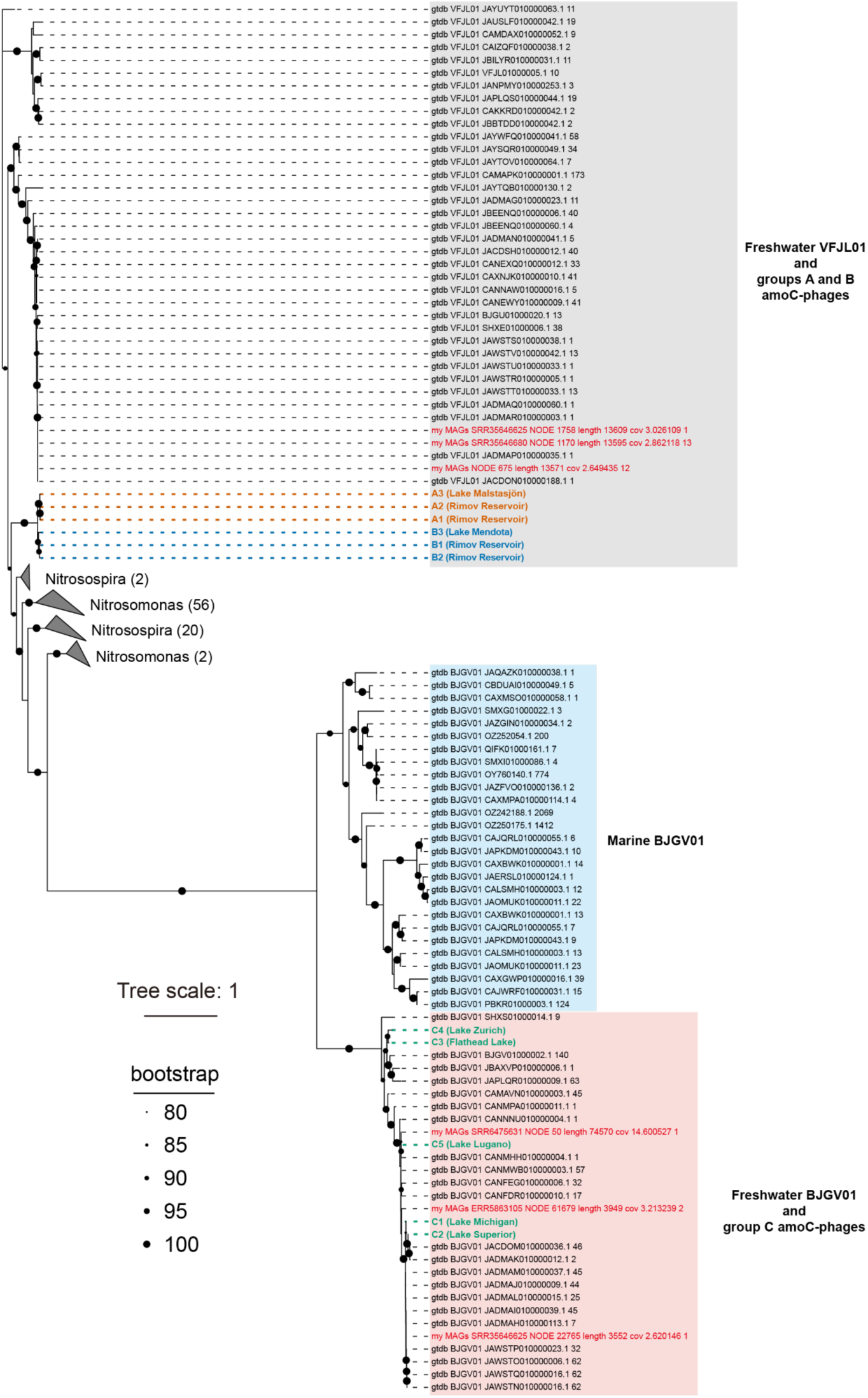
Phylogeny of phage- and bacteria-encoded phosphate-selective porin genes (*oprP*). Phylogenetic placement of *oprP* genes encoded by amoC-phages, reconstructed VFJL01 and BJGV01 genomes, and bacterial reference genomes (including all public VFJL01 and BJGV01 genomes). The tree includes OprP homologs from GTDB r226 genomes with >60% protein identity to phage-encoded OprP sequences, *oprP* genes from VFJL01 and BJGV01 genomes reconstructed in this study, and *oprP* genes encoded by amoC-phages. VFJL01 and BJGV01 genomes reconstructed in this study are labeled in red. AmoC-phage-encoded *oprP* genes are colored according to phage group, and the lakes from which the amoC-phage genomes were reconstructed are shown in parentheses after genome IDs. OprP sequences from *Nitrosospira* and *Nitrosomonas* reference genomes are collapsed, with the number of included sequences shown in parentheses after each genus name. Dashed lines indicate the positions of amoC-phage-encoded *oprP* genes. The tree scale bar indicates amino acid substitutions per site.

**Extended Data Fig. 7.**
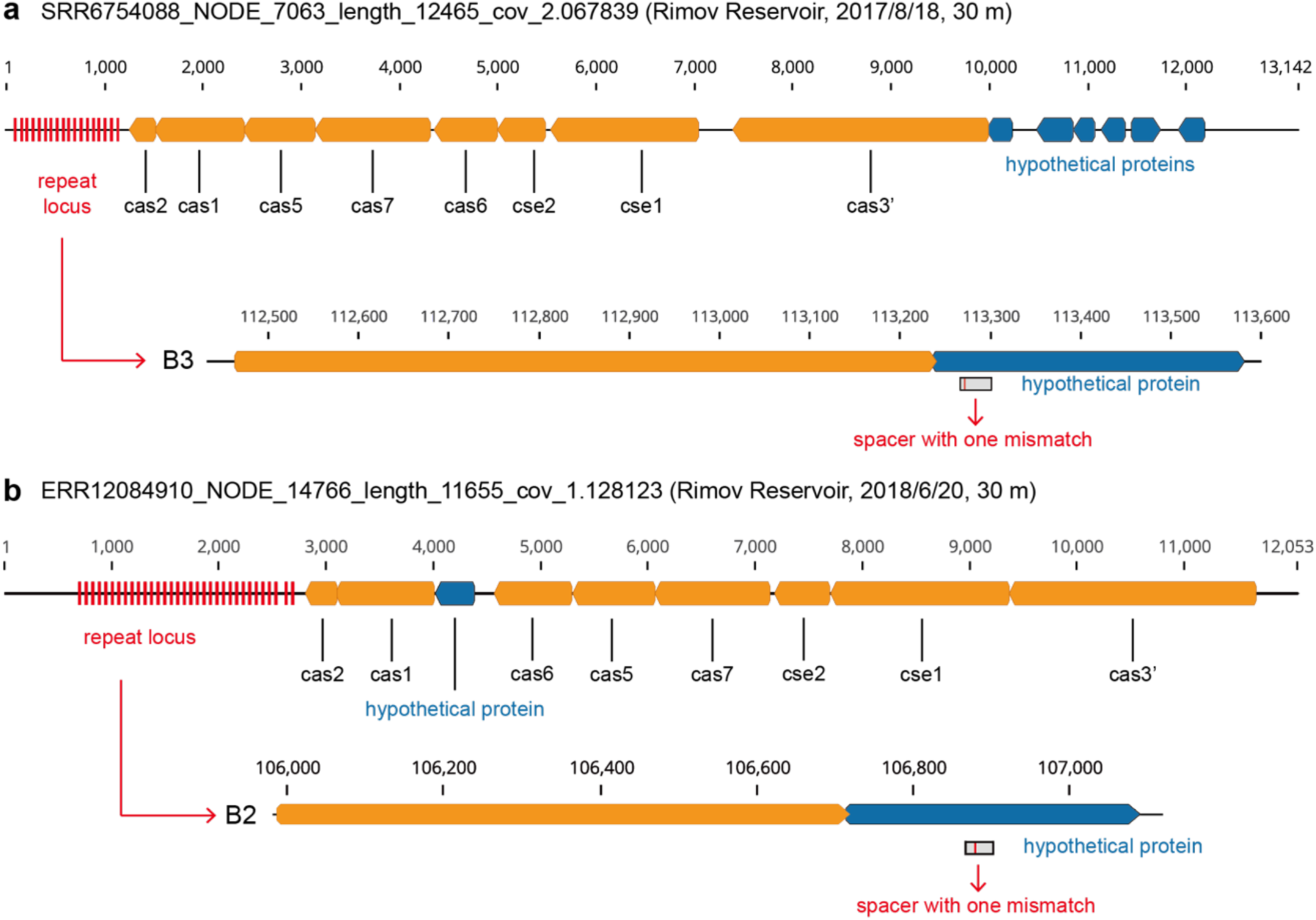
CRISPR spacers encoded by VFJL01 target group B amoC-phage genomes. Two VFJL01 scaffolds reconstructed from Rimov Reservoir metagenomes encode CRISPR-Cas loci with spacers targeting group B amoC-phages. (a) A spacer from a VFJL01 scaffold targets the B3 amoC-phage genome. (b) A spacer from a second VFJL01 scaffold targets the B2 amoC-phage genome. For each panel, the CRISPR-encoding VFJL01 scaffold is shown above, with the repeat locus and adjacent cas genes annotated, and the corresponding targeting region in the amoC-phage genome is shown below. Red arrows indicate the locations of the repeat loci. Both spacers contain one mismatch relative to the phage sequences. Scaffold identifiers, sampling dates, and depths are indicated above each CRISPR-encoding scaffold.

**Extended Data Fig. 8.**
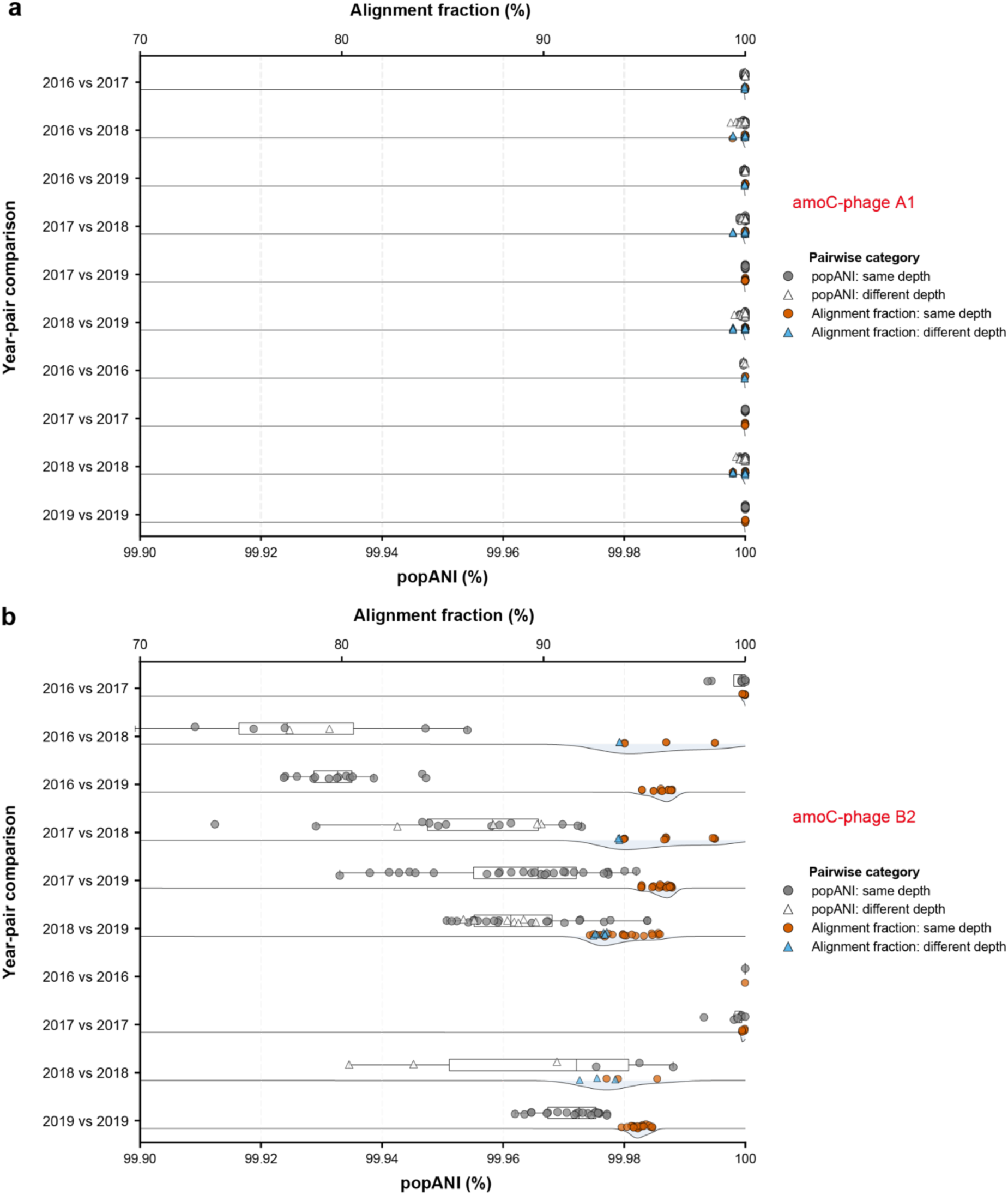
Population dynamics and structure of group A and B amoC-phages in Rimov Reservoir. (a, b) Pairwise population comparisons for amoC-phage A1 (a) and amoC-phage B2 (b) across Rimov Reservoir metagenomic samples. For each year-pair comparison, the lower x-axis shows pairwise population average nucleotide identity (popANI), and the upper x-axis shows pairwise alignment fraction. Boxplots summarize the distributions of popANI values, whereas ridge plots show the corresponding distributions of alignment fractions. Individual pairwise comparisons are overlaid and categorized according to whether the paired samples were collected from the same depth or from different depths. A1 showed consistently high popANI and alignment fraction across comparisons, indicating a genetically stable population through time. By contrast, B2 showed greater variation in both popANI and alignment fraction, indicating stronger temporal population structuring and turnover among closely related group B amoC-phage populations.

**Extended Data Fig. 9.**
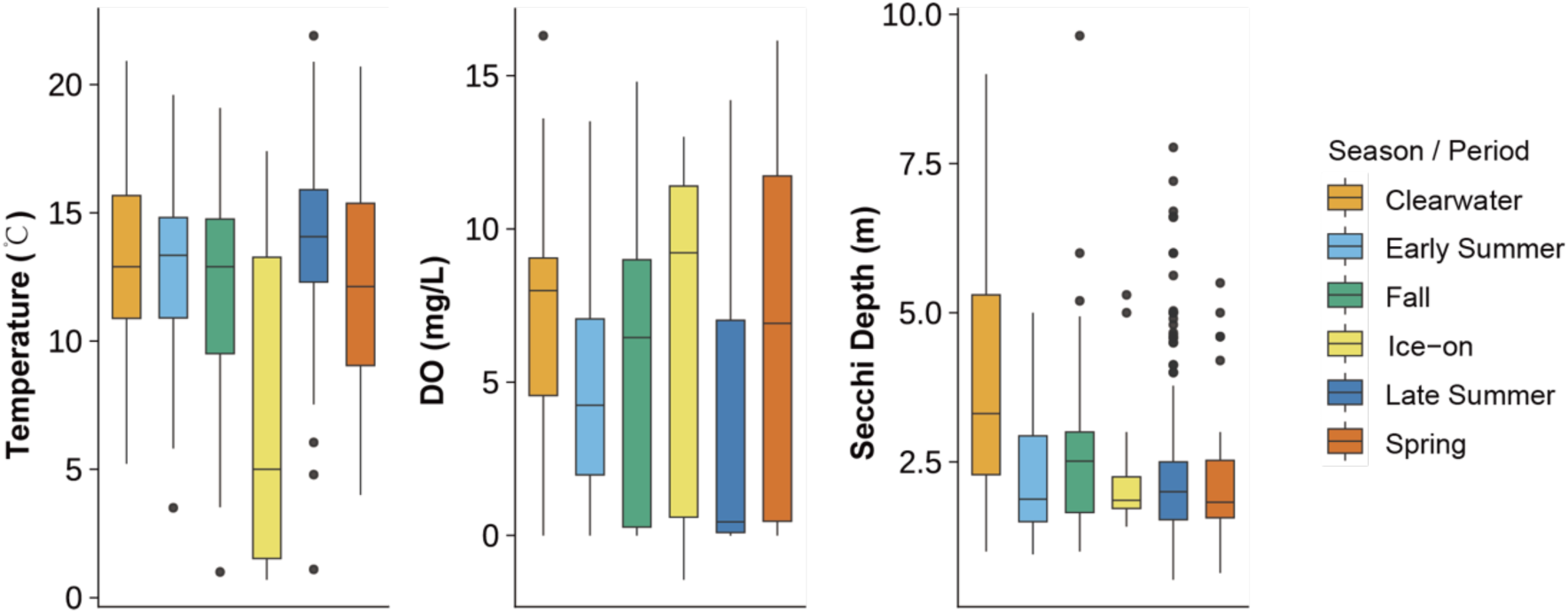
Physicochemical characteristics of Lake Mendota samples (2000-2019) across seasonal periods. Boxplots showing the distribution of temperature, dissolved oxygen (DO) concentration, and Secchi depth for the long-term Lake Mendota metagenomic samples analyzed in this study. Samples were grouped by seasonal or limnological period, including Clearwater, Early Summer, Fall, Ice-on, Late Summer, and Spring. Boxes indicate the interquartile range, horizontal lines indicate medians, whiskers extend to 1.5× the interquartile range, and points indicate outliers. The Ice-on samples were characterized by lower temperatures and relatively high dissolved oxygen concentrations compared with most open-water periods, whereas Secchi depth varied across seasons. Physicochemical metadata were obtained from the long-term Lake Mendota dataset reported in Zhou et al. ^18^.

**Extended Data Fig. 10.**
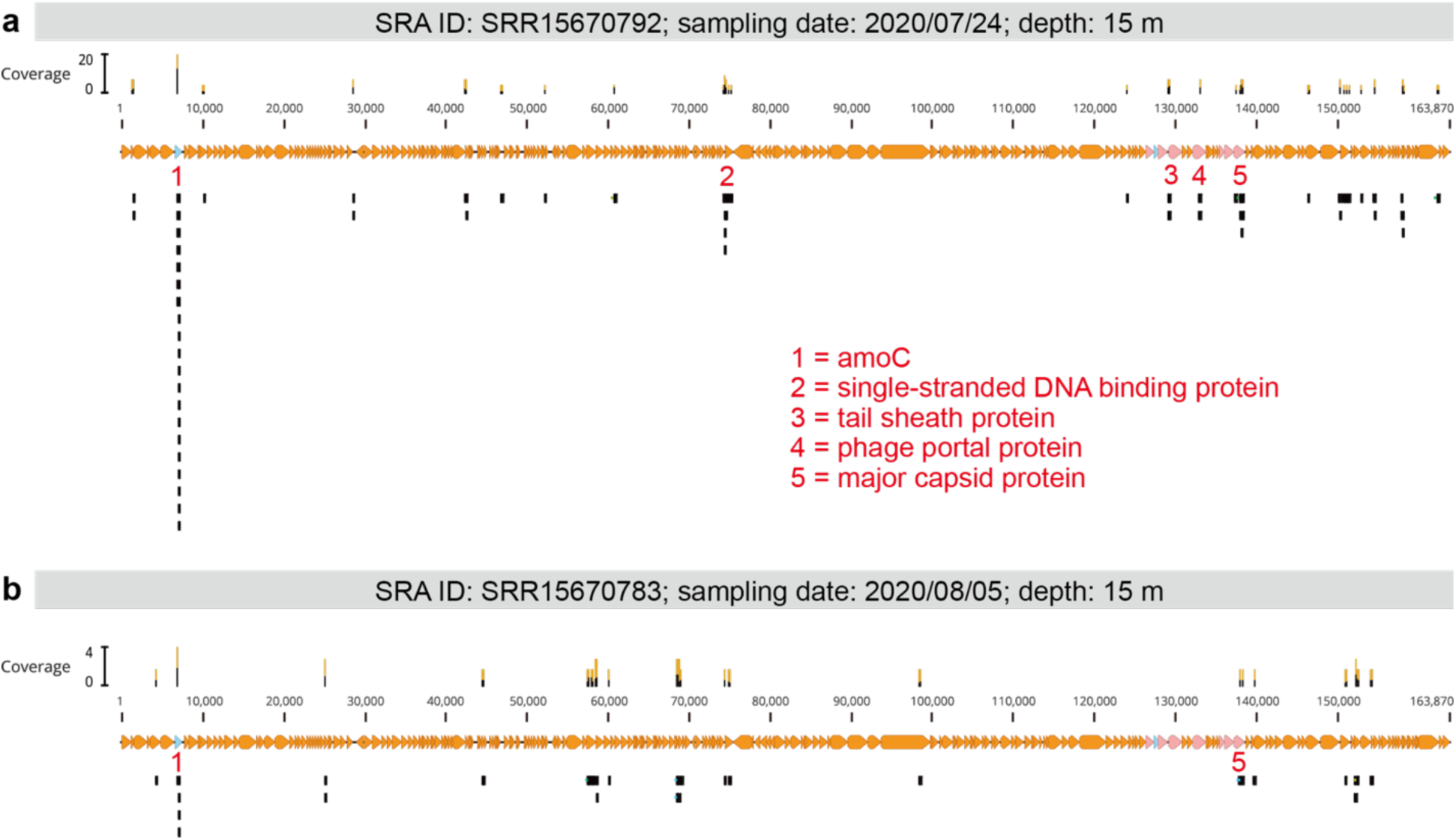
Limited transcriptional activity of group B amoC-phage B3 in Lake Mendota metatranscriptomes. Quality-filtered RNA reads from two 15 m Lake Mendota samples collected during summer 2020 were mapped to the B3 amoC-phage genome, allowing no more than 1% mismatches per read, and visualized in Geneious Prime. Coverage profiles of mapped RNA reads are shown above each genome map, and individual mapped RNA reads are shown as black blocks below the genes. Genes with detectable transcriptional signals are indicated by red numbers, with corresponding annotations shown in the center of the figure. Sample metadata, including SRA accession, sampling date, and depth, are shown in the grey bars above each panel. The sparse read coverage indicates limited but detectable transcriptional activity of the group B amoC-phage B3 genome in these two samples.

