## Supplementary Information for "Widespread phages exhibit depth-structured infection coupled with ammonia oxidation"

**for**

### Supplementary Notes S1: Host prediction based on AMGs

All amoC-phages except C3 also encode a phosphate-selective porin gene, *oprP* (Fig. 2c). Homology searches and phylogeny showed that phage-encoded OprP proteins grouped with homologs from their predicted freshwater hosts, VFJL01 or BJGV01, broadly mirroring the amoC-based host assignments (Extended Data Fig. 6). Groups A and B phage OprP proteins shared 60–76% amino acid identity with VFJL01 homologs, whereas group C phage OprP proteins showed substantially higher similarity to freshwater BJGV01 homologs, with 81–100% amino acid identity. Moreover, the group C amoC-phages encode another three AMGs, which respectively encode cysteine desulfurase activator complex subunit SufB, hopanoid biosynthesis-associated radical SAM protein HpnH, and NapC/NirT-family multiheme cytochrome c. Protein sequence homolog search and phylogeny suggest that they originated from freshwater BJGV01 homologs (Supplementary Figs. 1-3)

### Supplementary Notes S2: Why have freshwater amoC-phages not been reported yet?

An important question raised by this study is why *amoC*-encoding viruses have been previously reported in marine systems but remained undetected in freshwater ecosystems. Several factors likely contribute to this gap. First, marine and freshwater systems differ fundamentally in hydrodynamics, nutrient regimes, stratification patterns, and oxygen gradients [1–4], which strongly structure the ecological niches of ammonia oxidizers and their associated viruses. The freshwater amoC-phages identified here are genomically distinct from previously reported marine *amoC*-carrying viruses, limiting the effectiveness of marine references for freshwater discovery. Second, the three phage groups described here exhibit markedly different population genetic structures. Group A is geographically restricted and exhibits extremely low microdiversity, whereas groups B and C show extensive intra-population variation, which likely fragments assemblies and obscures genome recovery in short-read datasets [5,6]. Third, in some cases, the high sequence similarity between phage-encoded *amoC* and host *amoC* complicates confident assignment of this gene to viral genomes without extensive curation [5]. Together with the historical focus on ammonia-oxidizing microorganisms rather than their viruses, these ecological and methodological factors likely explain why freshwater amoC-phages have remained overlooked.

### Supplementary Notes S3: The limitations of this study

This study has two major limitations that warrant further investigation. First, although the consistent association between amoC-phages and their predicted hosts is supported by multiple lines of evidence, neither VFJL01 nor BJGV01 has been obtained in pure culture. While transcriptional evidence supports active infection and *amoC* expression, the quantitative contribution of viral *amoC* to overall ammonia oxidation rates remains to be determined. Future work combining controlled experiments with *in situ* rate measurements will be necessary to assess the infection dynamics of amoC-phages and the functional impact of viral modulation on nitrification. Second, the high microdiversity observed in groups B and C likely leads to an underestimation of their true diversity and distribution in short-read metagenomic datasets, highlighting the need for long-read sequencing approaches to resolve population structure.

### Supplementary methods S1: Manual curation and genome extension across host-like regions

As described in the Methods section of the main text, the amoC-encoding viral scaffolds were manually curated and extended using read recruitment and scaffold-end overlap information. Because several auxiliary metabolic genes encoded by the group C amoC-phages showed very high nucleotide similarity to homologous genes in their predicted bacterial hosts, freshwater BJGV01, particular caution was taken when scaffold extension approached regions containing host-like genes, such as amoC, phosphate-selective porin (oprP), and NapC/NirT family cytochrome c. In such cases, short-read assembly could generate ambiguous graph structures in which viral and host-derived scaffolds shared nearly identical terminal sequences or gene regions.

For each candidate amoC-encoding viral scaffold, reads were mapped back to the scaffold, and terminal regions were inspected manually. Scaffold extension was performed using consensus sequences inferred from read overhangs at scaffold ends. The extended terminal sequences were then searched against all assembled scaffolds from the same sample using BLASTn to identify candidate scaffolds with compatible end-sequence overlaps. Candidate joins were evaluated based on the length and identity of the overlap, read support across the putative junction, sequencing coverage patterns, and the gene-content context of the joined scaffolds.

When the extension reached regions containing genes highly similar to bacterial homologs, the local sequencing coverage often increased markedly relative to the surrounding viral genomic regions if the corresponding bacteria had a high abundance in the sample. Such elevated coverage was interpreted as a potential indication of shared recruitment from both viral and host genomic copies, and usually, the assembly of short paired-end reads using the de Bruijn graph-based assemblers will break at such locations. In these cases, BLASTn searches using the extended scaffold ends sometimes identified multiple candidate scaffolds that could be joined through near-identical end-sequence overlaps. These candidate scaffolds included both scaffolds with viral genomic context and scaffolds with host-like genomic context.

To avoid erroneous incorporation of host genomic fragments into viral genomes, candidate extensions were assessed conservatively. Candidate scaffolds were prioritized for joining when they (1) showed coverage comparable to the amoC-encoding viral scaffold under curation, (2) were supported by consistent read-pair or read-overhang evidence across the junction, and (3) encoded viral hallmark genes, phage structural genes, or other genes embedded in a viral genomic context. In contrast, candidate scaffolds were not used for viral genome extension when they (1) showed coverage patterns more consistent with those of the host genome in the corresponding sample, (2) encoded predominantly bacterial housekeeping or host-context genes, or (3) lacked viral genomic features beyond the shared host-like gene region. When multiple candidate scaffolds satisfied the terminal-overlap criterion, the final extension path was selected only when the sequencing coverage and gene-content context jointly supported viral origin. Ambiguous joins were left unresolved rather than forced.

This procedure was particularly important for the group C amoC-phage genomes encoding auxiliary metabolic genes with high sequence similarity to bacterial homologs, as mentioned above. These regions were therefore treated as potential repeat-like or shared regions during manual curation. Genome extension across such loci required concordant support from terminal read overhangs, scaffold-end overlap, local coverage continuity, and viral genomic context. Scaffolds inferred to derive from host genomes were excluded from the curated viral genome sequences, even when they shared high-identity terminal overlaps with the extending viral scaffold. This conservative curation strategy reduced the risk of misassembly or artificial chimerism between viral and host genomic fragments while retaining extensions supported by multiple independent lines of evidence.

We acknowledge that this manual curation procedure requires substantial experience in interpreting read-mapping patterns, assembly continuity, and gene-context information. Nevertheless, this approach can be applied to similar studies, particularly when viral scaffolds contain genes that share extremely high sequence identity with their predicted hosts or with other members of the microbial community.

**Supplementary methods S2: Validation of host-like regions in group C amoC-phage genomes**

Three genes encoded by group C amoC-phage genomes, including *amoC*, the phosphate-selective porin gene *oprP*, the cysteine desulfurase activator complex subunit SufB gene, the hopanoid biosynthesis associated radical SAM protein HpnH gene, and the NapC/NirT-family multiheme cytochrome c gene, showed high sequence similarity to homologs in their predicted BJGV01 hosts. Following the manual curation and genome-extension procedures described above, we recovered three complete and two partial group C amoC-phage genomes. Here, we use representative genomes reconstructed from Lake Michigan and Lake Superior to illustrate how the genomic regions containing these host-like genes were validated.

As shown in Fig. 6c,d, several metagenomic samples contained detectable group C amoC-phages but no detectable predicted BJGV01 hosts. These included Lake Michigan station MI27M samples collected on 2 August 2019 at 38 m and 94 m, Lake Michigan station MI41M samples collected on 3 August 2019 at 2 m and 38 m, the Lake Superior station SU01M sample collected on 25 August 2019 at 30 m, and Lake Superior station SU17M samples collected on 22 August 2019 at 2 m and 29 m. These host-negative but phage-positive samples provided an opportunity to validate whether the host-like genes were genuinely embedded within the phage genomes rather than introduced by host-derived assembly artifacts during manual curation and genome extension.

Metagenomic reads from samples in which group C amoC-phages were detected but predicted BJGV01 hosts were not detected were mapped to the corresponding curated phage genomes. In the regions containing the host-like genes, paired-end reads continuously spanned the boundaries between these host-like genes and adjacent phage genomic regions (Supplementary Fig. 5). These read-pair connections support the placement of the host-like genes within the phage genomes and argue against their inclusion as host-derived assembly chimeras. Therefore, despite the high similarity of the phage-encoded host-like genes to BJGV01 homologs, they are confidently assigned to group C amoC-phage genomes.

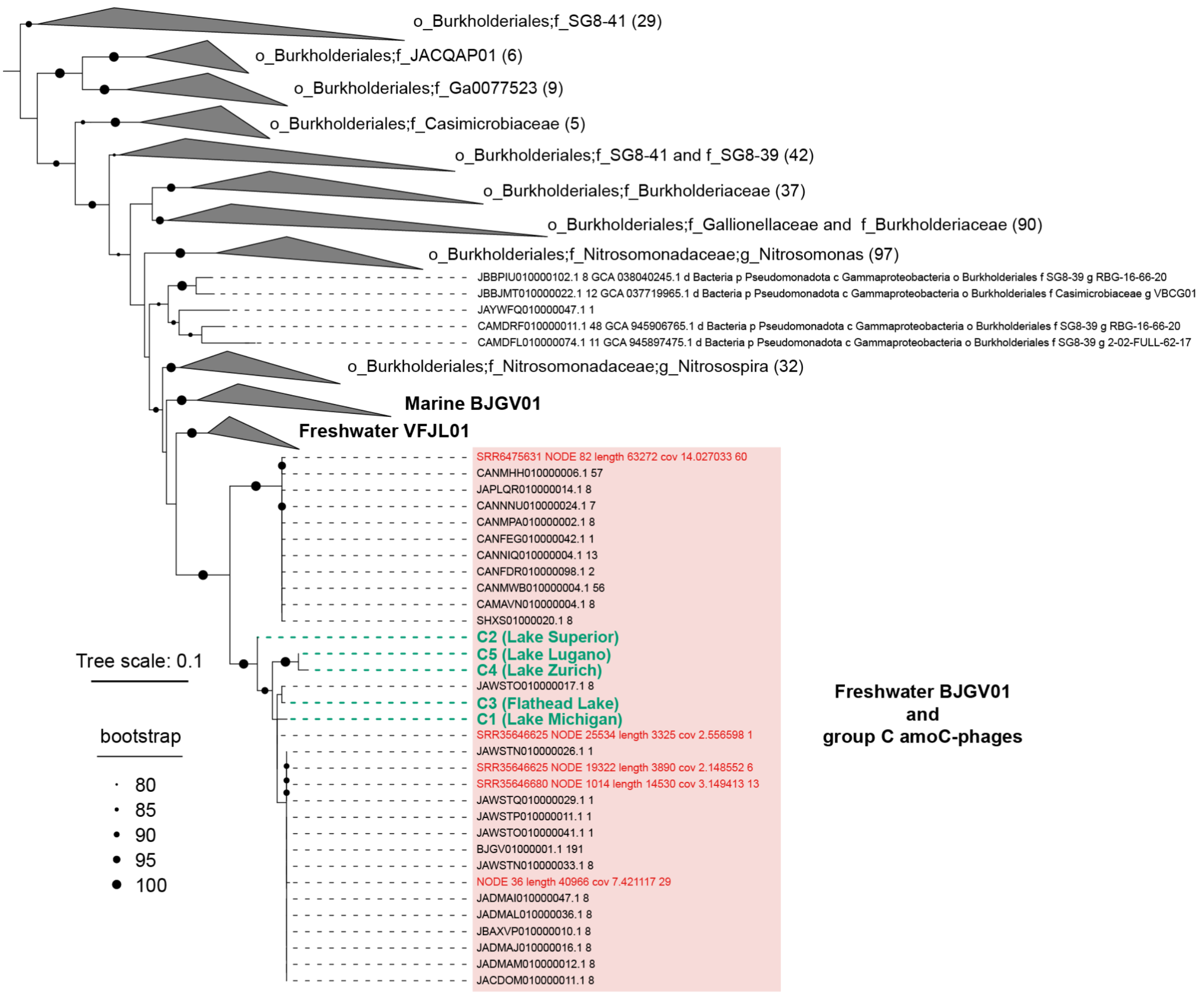


**Supplementary Fig. 1 | The phylogeny of phage- and bacteria-encoded SufB genes.** Phylogenetic placement of SufB genes encoded by the group C amoC-phages, reconstructed VFJL01 and BJGV01 genomes, and bacterial reference genomes (including all public VFJL01 and BJGV01 genomes). The tree includes SufB homologs from GTDB r226 genomes with >85% protein identity to phage-encoded sequences, SufB genes from VFJL01 and BJGV01 genomes reconstructed in this study, and SufB genes encoded by group C amoC-phages. VFJL01 and BJGV01 genomes reconstructed in this study are labeled in red. Group C amoC-phage-encoded SufB genes are labeled in green, and the lakes from which the amoC-phage genomes were reconstructed are shown in parentheses after genome IDs. SufB sequences from marine BJGV01, freshwater VFJL01, *Nitrosospira*, *Nitrosomonas,* and other reference genomes are collapsed, with the number of included sequences shown in parentheses after each family or genus name. Dashed lines indicate the positions of amoC-phage-encoded SufB genes. The tree scale bar indicates amino acid substitutions per site. The tree was rerooted using the f_SG8-41genes shown on the top.

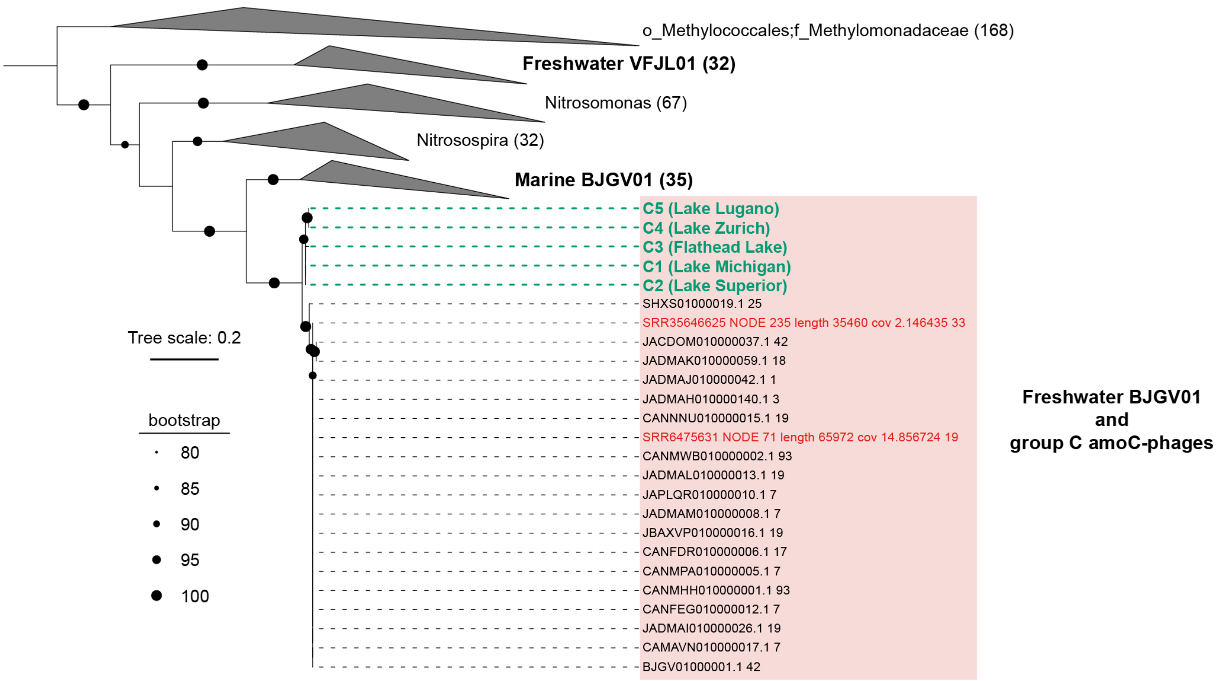


**Supplementary Fig. 2 | The phylogeny of phage- and bacteria-encoded HpnH genes.** Phylogenetic placement of HpnH genes encoded by the group C amoC-phages, reconstructed VFJL01 and BJGV01 genomes, and bacterial reference genomes (including all public VFJL01 and BJGV01 genomes). The tree includes HpnH homologs from GTDB r226 genomes with >80% protein identity to phage-encoded sequences, HpnH genes from VFJL01 and BJGV01 genomes reconstructed in this study, and HpnH genes encoded by group C amoC-phages. VFJL01 and BJGV01 genomes reconstructed in this study are labeled in red. Group C amoC-phage-encoded HpnH genes are labeled in green, and the lakes from which the amoC-phage genomes were reconstructed are shown in parentheses after genome IDs. HpnH sequences from marine BJGV01, freshwater VFJL01, *Nitrosospira*, *Nitrosomonas,* and other reference genomes (Methylomonadaceae) are collapsed, with the number of included sequences shown in parentheses after each family or genus name. Dashed lines indicate the positions of amoC-phage-encoded HpnH genes. The tree scale bar indicates amino acid substitutions per site. The tree was rerooted using the Methylomonadaceae genes shown at the top.

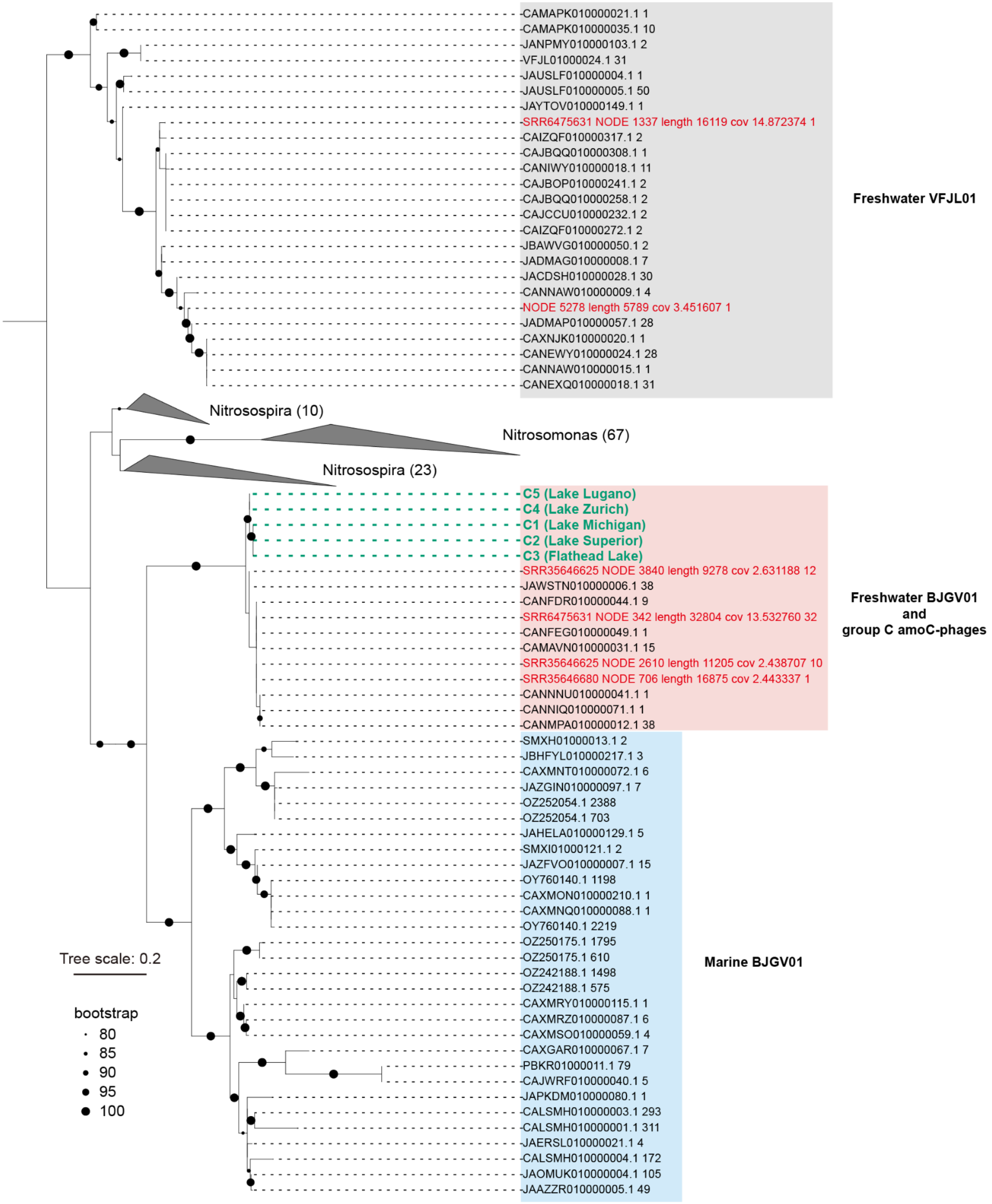


**Supplementary Fig. 3 | The phylogeny of phage- and bacteria-encoded NapC/NirT-family multiheme cytochrome c genes.** Phylogenetic placement of NapC/NirT-family multiheme cytochrome c genes encoded by the group C amoC-phages, reconstructed VFJL01 and BJGV01 genomes, and bacterial reference genomes (including all public VFJL01 and BJGV01 genomes). The tree includes NapC/NirT homologs from GTDB r226 genomes with >75% protein identity to phage-encoded sequences, NapC/NirT genes from VFJL01 and BJGV01 genomes reconstructed in this study, and NapC/NirT genes encoded by group C amoC-phages. VFJL01 and BJGV01 genomes reconstructed in this study are labeled in red. Group C amoC-phage-encoded NapC/NirT genes are labeled in green, and the lakes from which the amoC-phage genomes were reconstructed are shown in parentheses after genome IDs. NapC/NirT sequences from *Nitrosospira* and *Nitrosomonas* reference genomes are collapsed, with the number of included sequences shown in parentheses after each genus name. Dashed lines indicate the positions of amoC-phage-encoded NapC/NirT genes. The tree scale bar indicates amino acid substitutions per site. The tree was rerooted using the freshwater VFJL01 genes.

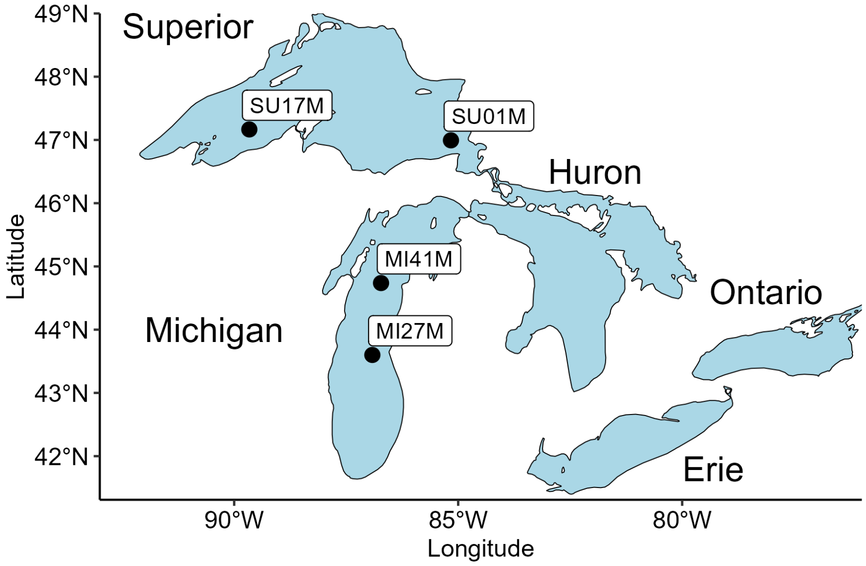


**Supplementary Fig. 4 | A brief map showing the locations of metagenomic and metatranscriptomic samples in Lake Michigan and Lake Superior in 2019.** Two locations in Lake Michigan and two locations in Lake Superior were included.

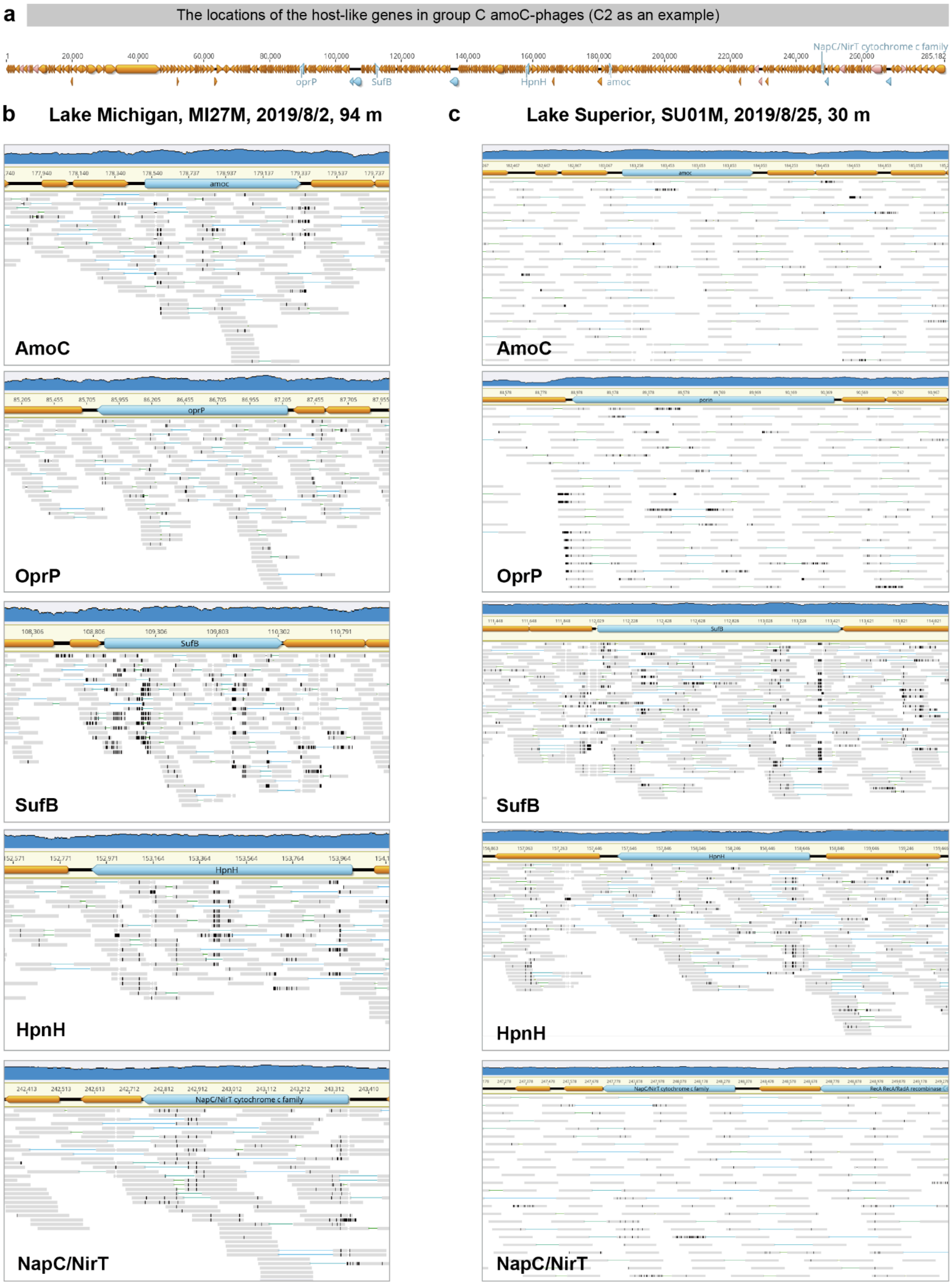


**Supplementary Fig. 5 | Read-mapping validation of host-like genomic regions in group C amoC-phage genomes.** (a) The locations of the host-like genes on the group C amoC-phages. The C2 genome reconstructed from Lake Superior is used as an example. Representative regions containing *amoC*, the phosphate-selective porin gene (*oprP*), the cysteine desulfurase activator complex subunit SufB gene, the hopanoid biosynthesis associated radical SAM protein HpnH gene, and the NapC/NirT-family multiheme cytochrome c gene are shown for group C amoC-phage genomes reconstructed from (b) a Lake Michigan sample and (c) a Lake Superior sample. Metagenomic read mapping across group C amoC-phage genomic regions encoding genes highly similar to those of their predicted BJGV01 hosts. Metagenomic reads from samples in which group C amoC-phages were detected, but predicted BJGV01 hosts were not detected, were mapped back to the curated phage genomes. Coverage profiles are shown above each genomic region, and mapped paired-end reads are shown below. Paired-end reads spanning the boundaries between the host-like genes and adjacent phage genomic regions support the correct placement of these genes within the phage genomes. These read-pair connections indicate that the presence of *amoC*, *oprP*, and the NapC/NirT-family multiheme cytochrome c gene in group C amoC-phage genomes is unlikely to result from host-derived assembly chimeras. Reads were mapped to the genome using Bowtie2 v2.5.4 with default parameters [7] and visualized in Geneious Prime [8] (release version, 2025.1.2).
